# Revisiting the activity of two poly(vinyl chloride)- and polyethylene-degrading enzymes

**DOI:** 10.1101/2024.03.15.585159

**Authors:** Anton A. Stepnov, Esteban Lopez-Tavera, Ross Klauer, Clarissa L. Lincoln, Ravindra R. Chowreddy, Gregg T. Beckham, Vincent G. H. Eijsink, Kevin Solomon, Mark Blenner, Gustav Vaaje-Kolstad

## Abstract

Biocatalytic degradation of non-hydrolyzable plastics is a rapidly growing field of research, driven by the global accumulation of waste. Enzymes capable of cleaving the carbon-carbon bonds in synthetic polymers are highly sought-after as they may provide tools for environmentally friendly plastic recycling. Despite some reports of oxidative enzymes acting on non-hydrolyzable plastics, including polyethylene or poly(vinyl chloride), the notion that these materials are susceptible to efficient enzymatic degradation remains controversial, partly driven by a general lack of studies independently reproducing previous observations. We attempted to replicate two recent studies reporting that deconstruction of polyethylene and poly(vinyl chloride) can be achieved using an insect hexamerin from *Galleria mellonella* (so-called “Ceres”) or a bacterial catalase-peroxidase from *Klebsiella sp.*, respectively. Reproducing previously described experiments with the recombinant proteins, we did not observe any activity on plastics using multiple reaction conditions and multiple substrate types. Digging deeper into the discrepancies between the previous data and our observations, we show how and why the original experimental results may have been misinterpreted, leading to the erroneous claim that enzymatic deconstruction of polyethylene and poly(vinyl chloride) had occurred. Our results should lead to caution when interpreting the growing amount of literature claiming enzymatic degradation of non-hydrolyzable plastics.

## Introduction

Biocatalysis may represent a viable strategy to enable the deconstruction of recalcitrant polymer waste, as illustrated by a wide-spread use of hydrolytic and oxidative enzymes in industrial depolymerization of plant biomass [1]. The hydrophobicity and crystallinity of lignocellulose in plant cell walls resemble the properties of many types of plastics [2], so it is reasonable to believe that enzymatic conversion of synthetic polymers occurs in nature or can be achieved by enzyme engineering. Indeed, poly(ethylene terephthalate) (PET) is enzymatically depolymerized through ester hydrolysis in the polymer backbone. The first PET-active enzyme was discovered in 2005 [3], creating a new and promising field of research. Encouragingly, effective enzymatic deconstruction strategies for PET have been developed such that full depolymerization of this synthetic polyester can be achieved [4].

At the same time, many types of modern commodity plastics feature non-hydrolyzable carbon-carbon bonds in their backbones (e.g., polyethylene, polypropylene, polystyrene, and (poly)vinyl chloride) [4], all of which are highly resistant to depolymerization. Mechanisms for the biodegradation of non-hydrolyzable plastics are poorly understood and technologies for biocatalytic conversion are not yet available. Most research considering biodegradation of non-hydrolyzable plastics has focused on polyethylene (PE) [5]. Some (varying) degree of PE degradation by bacteria, fungi, and insects has been reported in the past [6–11]. On the protein level, such activity has typically been attributed to various oxidative enzymes, such as laccases and peroxidases [12–15]. Despite these claims, the notion that PE can be effectively degraded in Nature by enzymes is met with scepticism [4, 16]. The interpretation of existing data on biotic deconstruction and metabolization of PE is complicated by the fact that many studies rely on relatively non-precise analytical methods, such as gravimetry, microscopy, or infrared spectroscopy. Another complication lies in the use of plastics that are not pure and that may contain metabolizable additives. As pointed out by Wei *et al*. [16], isotope tracing experiments can provide strong evidence for the biotic conversion of PE and other types of non-hydrolyzable plastics. Such studies are rare, and they report exceedingly slow PE metabolization rates that often tend to level off quickly [17, 18]. While these observations may be taken to indicate that some organisms indeed have the enzymes to convert PE, the slow rates and the limited extent of the processes may also indicate that only a minor, less recalcitrant fraction of the material is metabolized. When critically evaluating existing data, the potential role of abiotic disruption of the polymer surface by photo-oxidation should also be considered [5, 19].

From a biochemical perspective, it cannot be excluded that some organisms can indeed degrade and metabolize PE. Long chain linear alkanes, which can be viewed as low molecular weight PE mimics, are well-known to be used by some bacteria as a carbon source. Alkane catabolism is achieved by the consecutive action of oxidative enzymes, ultimately converting alkanes to corresponding bio-accessible fatty acids [20, 21]. Therefore, metabolism of PE and, perhaps, other plastics with a backbone of saturated carbon-carbon bonds, could be envisaged if there were to exist enzymes capable of converting the polymer to shorter fragments that could then enter routes for alkane biotransformation. Such enzymes would need to exhibit a unique ability to activate high-energy carbon-carbon bonds in insoluble and very hydrophobic substrates and would be of immense fundamental and applied value.

Encouragingly, two recent publications reported that oxidative degradation of non-hydrolysable plastics can be achieved at mild conditions using novel redox enzymes. Zhang *et al*. claimed that a recombinantly produced bacterial catalase-peroxidase from *Klebsiella sp.* EMBL-1 is capable of deconstructing poly(vinyl chloride) (PVC) [22]. Similarly, Sanluis-Verdes *et al*. suggested that two recombinantly produced hexamerins found in the saliva of greater wax moth larvae (*Galleria mellonella*), called “Ceres” and “Demetra” by the authors, catalyze deconstruction of PE [23]. Importantly, in contrast to other publications, the studies by Zhang *et al*. and Sanluis-Verdes *et al*. feature advanced and direct analytical methods, such as size exclusion chromatography (SEC), to assess the molar mass distribution in plastics, and gas chromatography-mass spectrometry (GC-MS) to detect cleavage products.

Intrigued by these findings, we have produced the bacterial catalase-peroxidase and the protein called “Ceres”, using expression approaches comparable to the ones used in the original reports, and carried out an extensive analysis of their activity towards PVC and PE, respectively. After testing multiple reaction conditions and substrates, no degradation or oxidation of plastics was observed, despite the use of multiple independent analytical methods including pyrolysis-GC-MS and SEC. Furthermore, we were unable to detect deconstruction of PE film treated with crude saliva from *Galleria mellonella* larvae, in contrast to previous reports. Digging deeper into discrepancies between the previously published data and our own observations, we were able to show how and why some of the original results from these studies were misinterpreted. While we cannot rule out the existence of biological deconstruction of non-hydrolyzable plastics, our study provides a cautionary outlook on the growing amount of literature claiming enzymatic degradation of such polymers and reveals potential reasons behind non-reproducibility and misinterpretation of experimental data.

## Results

### Catalase-peroxidase from *Klebsiella* sp. EMBL-1 is an active enzyme capable of oxidizing Amplex Red

The catalase-peroxidase from *Klebsiella* sp. EMBL-1 (referred to as *Kleb*CP hereafter) was produced in *E. coli* and purified by metal affinity chromatography (Fig. S1), yielding approximately 30 mg of the enzyme per 0.5 L of culture. The peroxidase activity of *Kleb*CP was confirmed using Amplex Red, a model substrate (Fig. 1). Note that such activity depends on H_2_O_2_ and that a 1:1 stoichiometry between H_2_O_2_ and the fluorescent reaction product (resorufin) is expected based on literature data on enzymatic peroxidation of Amplex Red [24]. In case of *Kleb*CP, consecutive addition of H_2_O_2_ (to 100 µM final concentration each time) showed that only a fraction of the H_2_O_2_ was used to oxidize Amplex Red. The progress curves indicate that even the third addition of 100 µM H_2_O_2_ leads to product formation, despite the total amount of Amplex Red in the reaction being only 100 µM. This confirms the bifunctional nature and the presence of catalase activity in *Kleb*CP leading to a non-productive H_2_O_2_ depletion (2H_2_O_2_ → O_2_ + 2H_2_O). Amplex Red oxidation by *Kleb*CP was not observed in the absence of the peroxidase co-substrate (H_2_O_2_), as expected.

**Figure 1.**
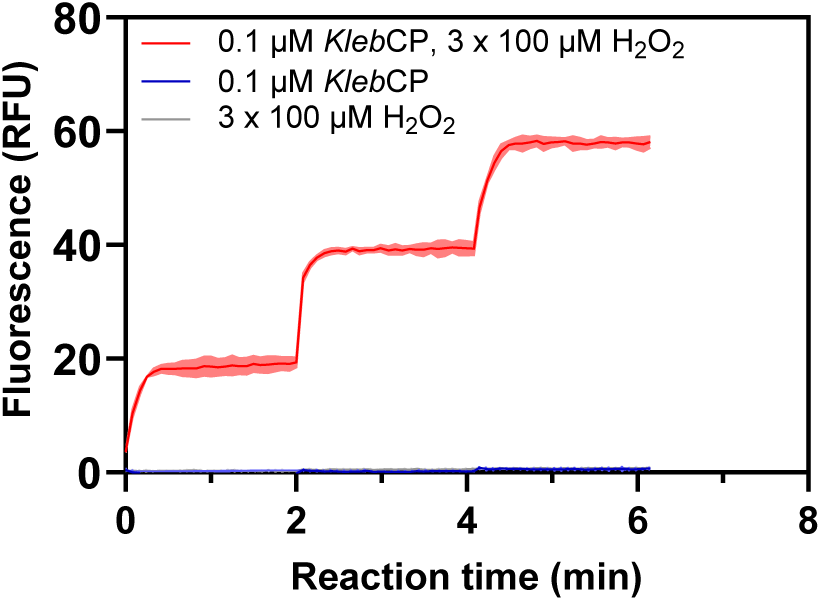
Peroxidase activity of *Kleb*CP on a small molecule substrate. The figure shows the formation of resorufin resulting from H_2_O_2_-dependent oxidation of 100 µM Amplex Red by 0.1 μM *Kleb*CP. The reactions were conducted at 30° C in 50 mM Tris-HCl buffer, pH 8.0, containing 300 mM NaCl. Note that H_2_O_2_ was introduced to the reaction mixtures 3 times at 0, 2 and 4 minutes of the experiment (each time to 100 µM final concentration). Also note that the 100 µM Amplex Red was not consumed after two additions of 100 µM H_2_O_2_, whereas the added H_2_O_2_ is fully depleted; this shows that a considerable fraction of the added H_2_O_2_ is removed by the catalase activity of *Kleb*CP. Error envelopes indicate standard deviation between replicates (*n*=3).

### *Kleb*CP does not cause oxidation or deconstruction of PVC

The capacity of *Kleb*CP to deconstruct PVC was then assessed, in an attempt to confirm the previous claims made by Zhang *et al*. [22]. In the prior study, incubation of 100 mg of PVC powder with 50 μg/mL (≈0.6 µM) *Kleb*CP for 96 hours at 30 °C resulted in an approximately 12% decrease in the number average molar mass (M_n_) of the polymer, which was assessed using SEC. In contrast, our SEC analysis did not show significant changes in the molar mass distribution after treating two different PVC powders (“high molecular weight” and “low molecular weight” PVC grades purchased from Sigma-Aldrich) with 10 µM *Kleb*CP using the same reaction conditions as previously reported (Fig. 2).

**Figure 2.**
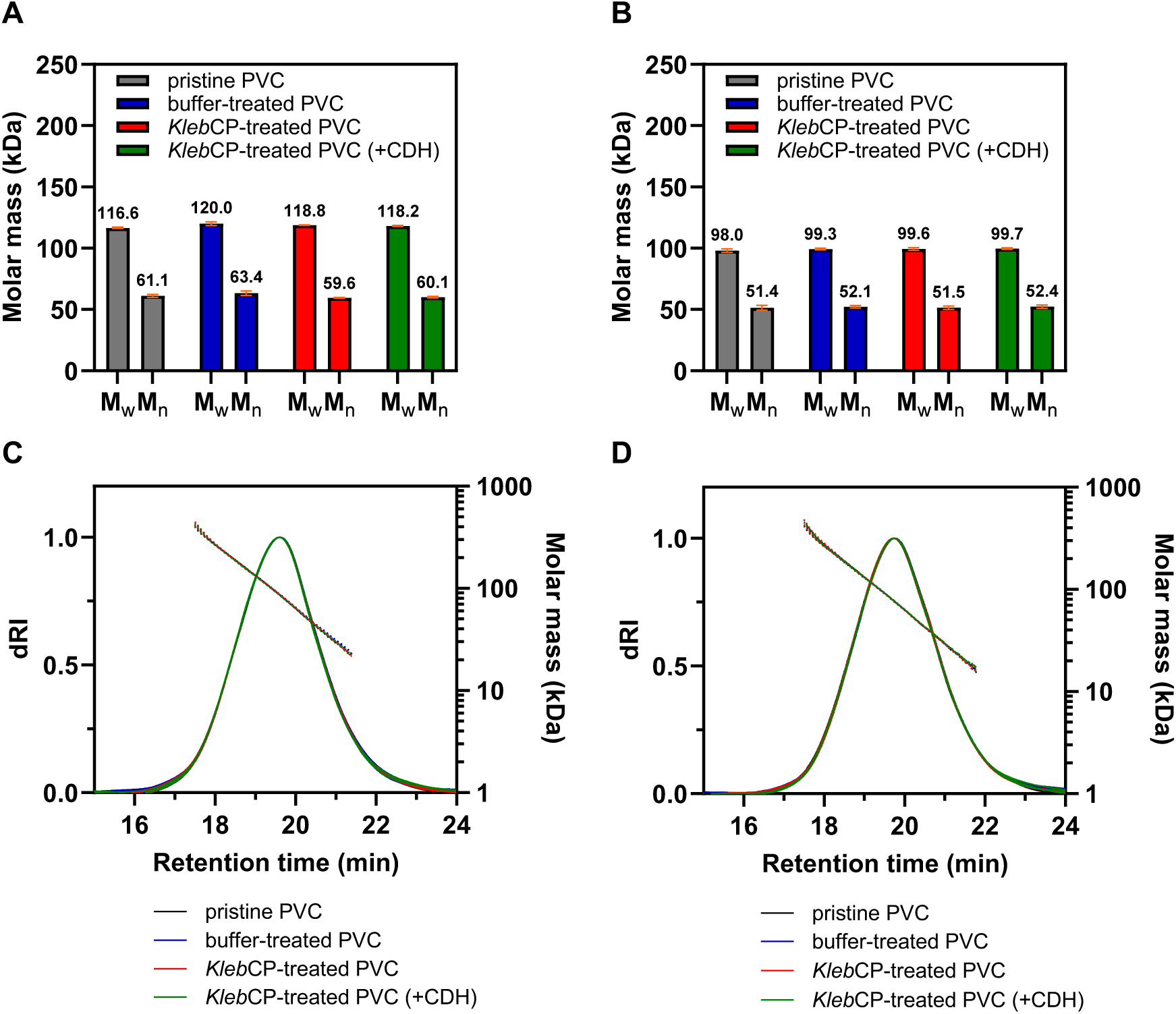
Effect of *Kleb*CP on the molar mass distribution in PVC samples. The figure shows the results of SEC analysis of high molecular weight (panels A, C) and low molecular weight (panels B, D) PVC samples treated with 10 µM *Kleb*CP in the absence or presence of an H_2_O_2_ source. 50 nM cellobiose dehydrogenase (CDH) was used alongside 5 mM cellobiose (CB) to generate hydrogen peroxide *in situ*. The reactions were incubated at 30 °C for 96 hours in 50 mM Tris-HCl buffer, pH 8.0, supplemented with 300 mM NaCl and 100 mg/mL PVC powder. The data featured in panels A and B were produced by averaging the results obtained for 3 independent reactions. Error bars indicate standard deviations between triplicates. M_w_, weight average molar mass; M_n_, number average molar mass. Panels C and D show chromatograms resulting from SEC-MALS (multi-angle light scattering) analysis of the samples. Solid lines indicate the differential refractive index detector response, whereas dotted lines represent the molar mass data. Note that the chromatographic traces are overlapping almost completely.

Both our experiments and the original experiments by Zhang *et al*. did not involve addition of the peroxidase co-substrate (H_2_O_2_). Given the lack of H_2_O_2_, it is not clear why a peroxidase reaction on plastic was observed in the original study. It cannot be excluded that some of the additives (e.g., antioxidants) known to be present in PVC [25] are capable of leaching into solution and reducing molecular oxygen, thus providing an *in situ* H_2_O_2_ source to fuel peroxidase reactions. To address this possibility, H_2_O_2_ production was monitored in the reaction mixtures in the presence of PVC (Fig. S2). This experiment indeed showed slow and delayed H_2_O_2_ generation starting to occur after approximately 8 h of incubation with both types of PVC substrates. To increase the level of H_2_O_2_ available to *Kleb*CP, we conducted PVC degradation experiments again, this time supplementing the reaction mixtures with 50 nM engineered cellobiose dehydrogenase (CDH) from *Crassicarpon hotsonii* [26] alongside 5 mM cellobiose. This variant of CDH is known to be capable of steady H_2_O_2_ generation by coupling cellobiose oxidation to the reduction of molecular oxygen, as shown in Fig. S2. Despite providing the *in situ* H_2_O_2_ source, the *Kleb*CP-treated plastics did not show any changes in the molar mass distribution (Fig. 2).

Looking for potential signs of PVC surface oxidation by the catalase-peroxidase, the enzyme-treated samples were further analyzed using FTIR spectroscopy. The treatment of PVC samples with *Kleb*CP resulted in formation of two weak peaks centred at approximately 1,530 cm^-1^ and 1,650 cm^-1^, accompanied by a broad peak located between 3,150-3,650 cm^-1^ (Fig. 3). These *Kleb*CP-associated signals are similar to the ones previously observed by Zhang *et al*., who interpreted two of these peaks as signs of double bond formation in the polymer backbone (1,530 cm^-1^) and hydroxylation (3,150-3,650 cm^-1^), i.e., signs of PVC oxidation [22]. However, deposition of *Kleb*CP on the surface of the FTIR sensor in the absence of PVC clearly showed that all the peaks putatively associated with PVC oxidation correspond to signals from the enzyme and not from PVC (Fig. 3). Furthermore, when *Kleb*CP-treated PVC was subjected to washing with Milli-Q water, the residual protein was removed from the PVC surface and the 1,530 cm^-1^ and 3,150-3,650 cm^-1^ signals disappeared (together with 1,650 cm^-1^ peak), consistent with these signals reflecting the enzyme (Fig. 3). Thus, the FTIR analysis clearly shows that the peaks previously claimed to indicate enzymatic oxidation of PVC result from adsorbed residual protein and not from PVC oxidation. Indeed, FTIR signals around 1,550 cm^-1^, 1,650 cm^-1^ and 3,000-3,500 cm^-1^ are well known to be observed in the presence of proteins [27, 28]. Overall, the results of our experiments with *Kleb*CP do not support its enzymatic activity on PVC.

**Figure 3.**
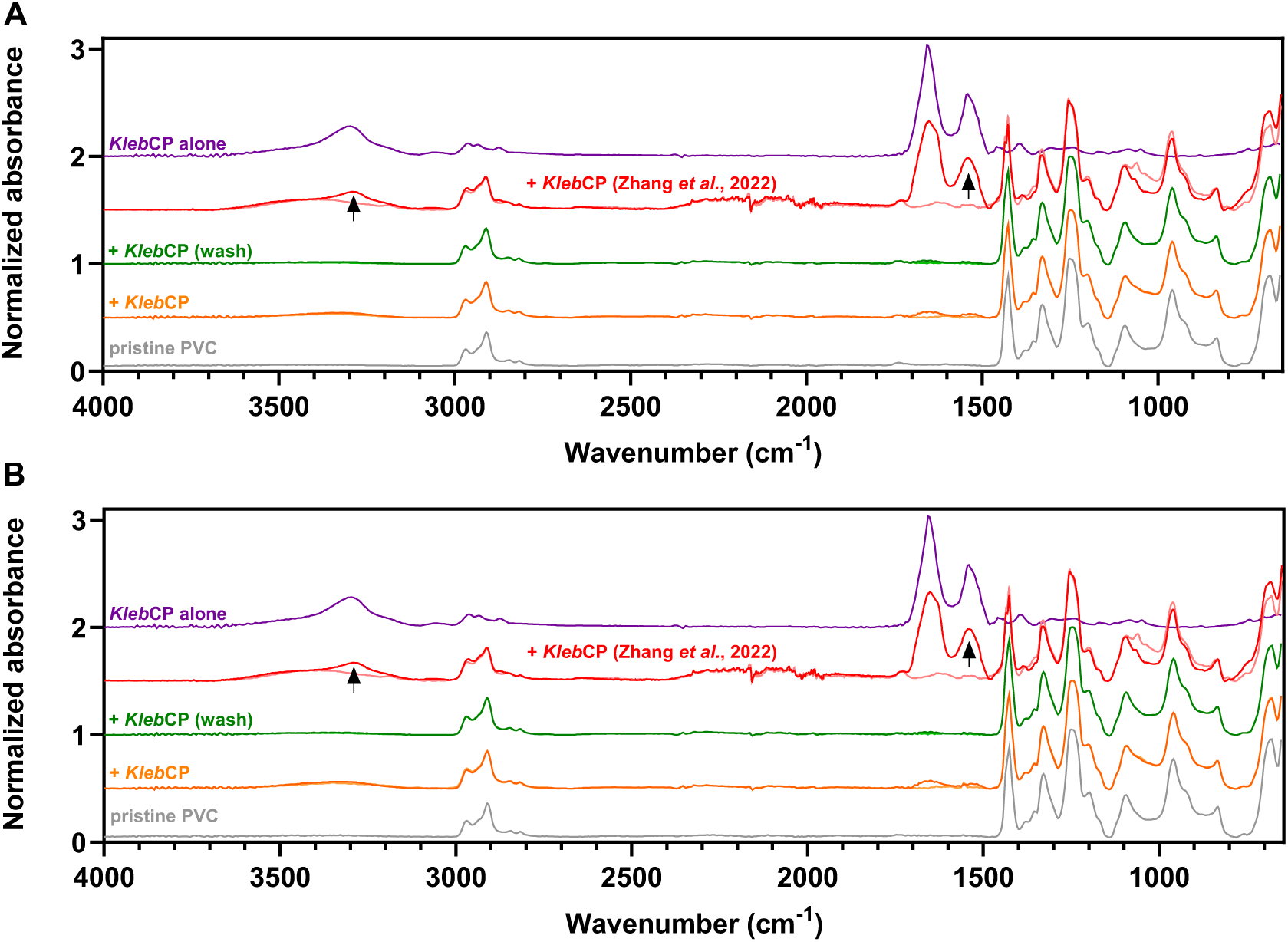
FTIR analysis of PVC samples after treatment with the catalase-peroxidase *Kleb*CP. The figure shows FTIR spectra of high molar mass (panel A) and low molar mass (panel B) PVC powders (100 mg/mL) after 96 hours of incubation at 30 °C with or without 10 µM *Kleb*CP in 50 mM Tris-HCl buffer, pH 8.0, containing 300 mM NaCl. Each FTIR trace represents the average of spectra obtained for 3 independently prepared reactions. The samples were first analyzed without removing the residual enzyme (orange trace) and then subjected to additional washing with Milli-Q water, as indicated in the figure (green trace). The light orange trace (superposed on the orange trace, no offset) represents FTIR spectra obtained for the control reactions lacking the enzyme. The FTIR data obtained for *Kleb*CP in the absence of any plastics is shown as a purple trace (“*Kleb*CP alone”), whereas the FTIR spectra of *Kleb*CP-treated PVC and buffer-treated PVC that were reported by Zhang *et al*. are shown in red and light red, respectively (superposed spectra; no offset between these two). These previously published spectra were plotted using the source data file provided with the original paper [22] and normalized as described below. Comparison of the traces clearly shows that the peaks previously claimed to indicate enzymatic oxidation of PVC (marked with black arrows) result from the adsorbed residual protein and not from true reaction products. The data were acquired with 8 cm^-1^ resolution. The absorbance signals were normalized using the C–H bend peak (≈ 1,250 cm^-1^) as the reference in PVC-containing samples and the amide I peak (≈ 1,650 cm^-1^) in the protein-only sample. Note that quantitative comparison between the data obtained for *Kleb*CP alone and for the samples containing PVC is not possible due to the difference in normalization.

### Cloning and production of “Ceres”

To gain insights into the previously described polyethylene degrading activity of hexamerins from the greater wax moth that were dubbed “Ceres” and “Demetra” by Sanluis-Verdes *et al*. [23], both proteins were initially targeted for recombinant production. However, our analysis of the protein sequences reported in the original study (NCBI accession numbers XP_026756459.1 and XP_026756396.1 for “Ceres” and “Demetra”, respectively), revealed that “Demetra” does not contain a detectable Sec/SPI signal peptide, in contrast to “Ceres” (Fig. S3). Given that such secretion signals are expected to be found in insect hexamerins [29], we took this observation as a sign that the sequence of the “Demetra” gene may be incomplete or inaccurate. Therefore, we focused our attention on “Ceres”. On a side note, a new version of the *Galleria mellonella* whole genome became available during the preparation of this manuscript (NCBI accession number NW_026442147.1), containing another variant of the “Demetra” gene (NCBI accession number XP_052756923.1). In contrast to the gene discussed by Sanluis-Verdes *et al*., the new gene variant encodes for a putative Sec/SPI signal peptide (Fig. S3), confirming our suspicion regarding the quality of the original “Demetra” sequence.

The synthesis of the “Ceres” gene and the recombinant protein production in Sf9 insect cells was carried out by Genscript, the same commercial provider as the one used in the original study by Sanluis-Verdes *et al*. [23]. Approximately 10 mg of electrophoretically pure hexamerin was produced from 2 L of culture (Fig. S4). The protein was then subjected to tryptic digestion, and the resulting protein fragments were analyzed using liquid chromatography with tandem mass spectrometry (LC-MS/MS) confirming that the protein in question indeed had the “Ceres” sequence (Fig. S5; 46% sequence coverage). Finally, the copper content of the protein was determined by inductively coupled plasma mass spectrometry (ICP-MS), to address the suggestion that “Ceres” may represent a copper-binding protein [30]. The amount of copper in the sample was found to be negligible, clearly showing that this protein does not bind copper (4.2 ± 2.3 nM Cu in 1,000 nM “Ceres” solution). Of note, the “Ceres” protein lacks all histidine residues that are known to bind copper in homologous phenoloxidases (Fig. S3; see below).

### “Ceres” is devoid of phenoloxidase activity

The first step after obtaining recombinant, pure “Ceres” was to test the protein for activity on a small molecule model substrate for phenoloxidases. According to the report of Sanluis-Verdes *et al*., hexamerins such as “Ceres” are phylogenetically related to phenoloxidases and may possess a similar type of activity. Phenoloxidases (POs) are copper-dependent enzymes that are ubiquitous in insects and able to oxidize L-3,4-dihydroxyphenylalanine (L-DOPA) to dopaquinone as a part of the melanin synthesis pathway [31]. L-DOPA oxidizing activity can indeed be detected in the haemolymph of *Galleria mellonella* [32]. In our work, the potential activity of “Ceres” on L-DOPA was assessed by monitoring both the formation of dopachrome (a product of the rapid auto-oxidation of dopaquinone) [33] and the depletion of molecular oxygen. Our results indicate that “Ceres” does not catalyze oxidation of L-DOPA (Fig. 4). To validate our experimental set up, tyrosinase from the common mushroom (*Agaricus bisporus*) was used as a positive control. This enzyme is known to possess a binuclear copper-site similar to the one found in insect POs and is capable of L-DOPA oxidation in the same manner [34]. Indeed, both the formation of dopachrome and the rapid depletion of O_2_ were detected when incubating the fungal tyrosinase with L-DOPA (Fig. 4).

**Figure 4.**
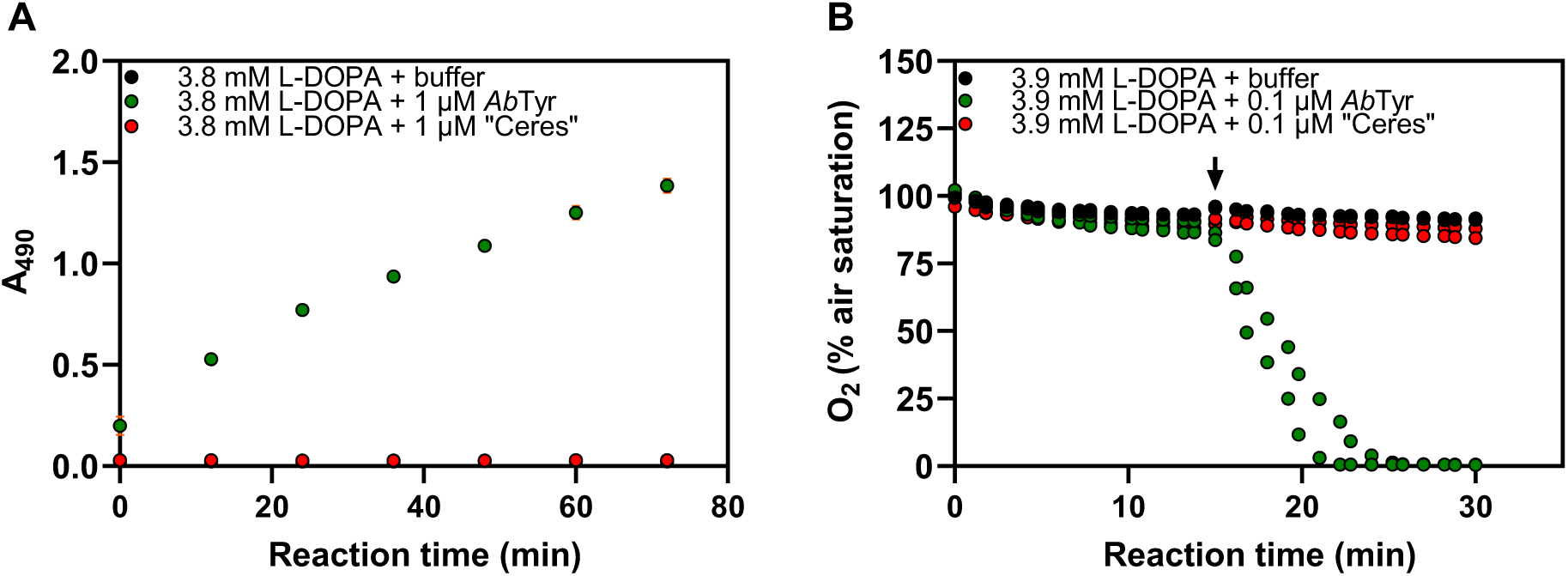
Probing the phenoloxidase activity of “Ceres” on L-DOPA. The figure shows the formation of dopachrome (panel A) and the depletion of molecular oxygen (panel B) as the result of L-DOPA oxidation by “Ceres” or by a tyrosinase from *Agaricus bisporus* (*Ab*Tyr). The reactions were carried out at room temperature in 20 mM HEPES, pH 7.0, supplied with 150 mM NaCl and 5% (v/v) glycerol, using various concentrations of enzymes and L-DOPA. Error bars in panel A represent standard deviations for triplicates and in most cases are hidden behind the data markers. The oxygen depletion experiments (panel B) were initiated by injecting enzymes or reaction buffer (negative control) through self-sealing septa into air-tight reaction vessels containing the substrate. The injections occurred at approximately 15 minutes of incubation, as indicated by the black arrow. The reactions shown in panel B were performed twice and all data points are shown.

### “Ceres” is not capable of oxidizing and deconstructing polyethylene films

To probe the capacity of “Ceres” to oxidize polyethylene films, LDPE film discs weighing ≈ 1.7 mg each were fully submerged in the reaction buffer described by Sanluis-Verdes *et al*. (20 mM HEPES, pH 7.0, supplied with 150 mM NaCl and 5% (v/v) glycerol), containing 1 mg/mL (≈ 12 μM) “Ceres”. The films were incubated for 70 h at room temperature, washed once with milli-Q water and once with 96% (v/v) ethanol, and subjected to FTIR analysis (Fig. 5A). We did not observe the formation of a ≈ 1,745 cm^-1^ (carbonyl) peak, previously reported by Sanluis-Verdes *et al*. as evidence for “Ceres”-dependent oxidation of PE. Meanwhile, the incubation of plastic films with “Ceres” resulted in the appearance of two new peaks in the ≈ 1,655 cm^-1^ and ≈ 1,545 cm^-1^ regions, which is a clear and well recognized sign of protein contamination [27, 28]. The same results (Fig. S6) were obtained when applying 5 μL drops of a 1 mg/mL “Ceres” solution in reaction buffer onto an LDPE film over 3 days (14 drops in total), which was similar to the approach originally used by Sanluis-Verdes *et al*. To ensure that the observed signal from the adsorbed residual protein did not interfere with the detection of potential PE oxidation, the “Ceres”-treated LDPE film discs were subjected to additional washing with 1 M NaOH. As expected, the signals that we believe are protein-related decreased significantly (Fig. 5B). No new peaks indicative of film oxidation were observed.

**Figure 5.**
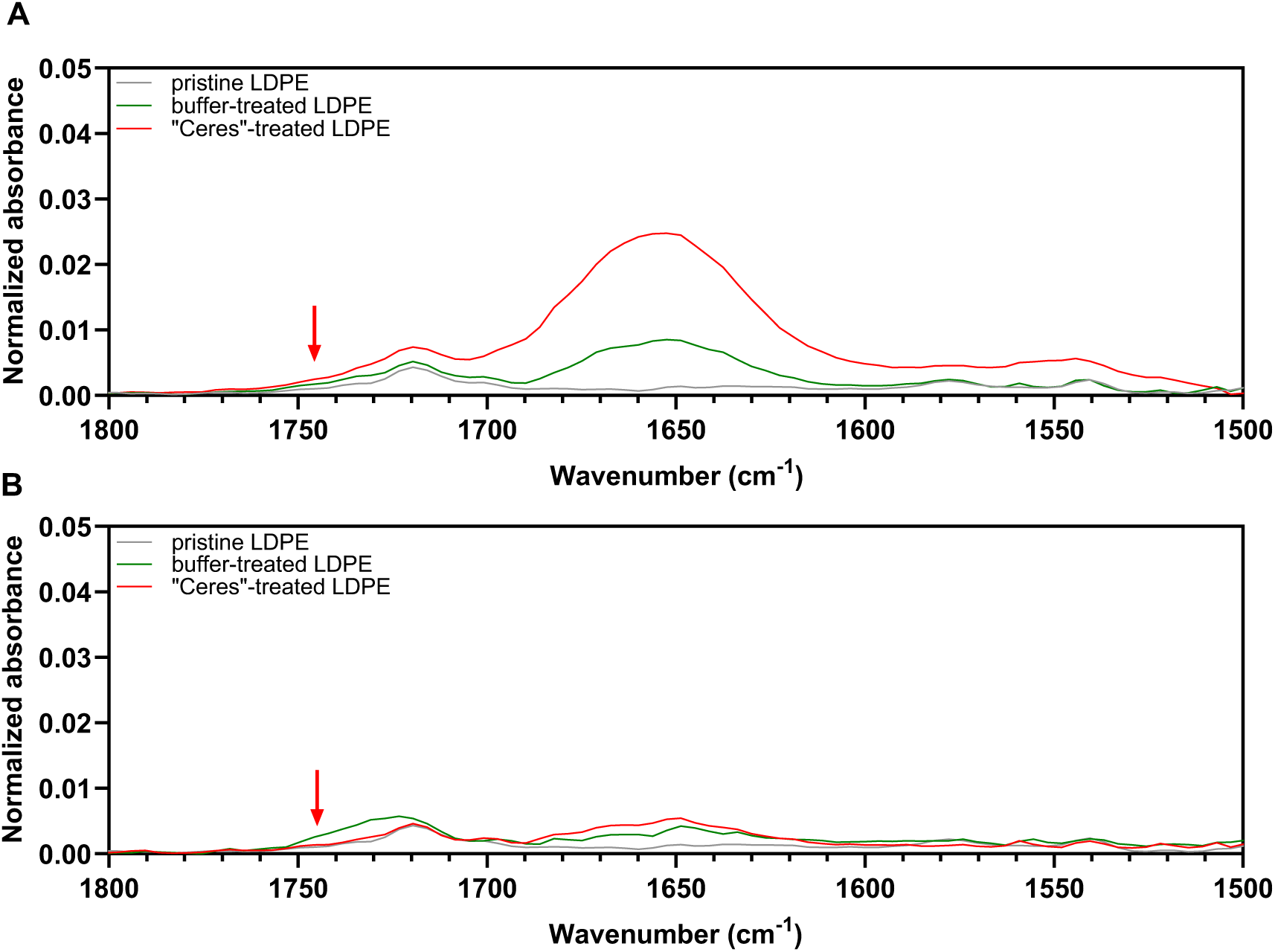
FTIR analysis of LDPE films treated with “Ceres”. The figure shows parts of the FTIR spectra of “Ceres”-treated films, buffer-treated films, and pristine non-treated films before (panel A) and after cleaning with 1 M NaOH (panel B). Red arrows indicate the expected position (≈ 1,745 cm^-1^) of a carbonyl peak which was not observed in our experiments but was previously reported as a result of polyethene oxidation by “Ceres”. Each trace shown in the figure represents the average of signals obtained for 3 film discs. The FTIR peaks resulting from the incubation of polyethene with “Ceres” that are visible in panel A are signals from the protein, and not from reaction products (see the main text for discussion). In-house made additive-free LDPE films were incubated at 25 °C for 70 hours in 20 mM HEPES, pH 7.0, supplied with 150 mM NaCl and 5% (v/v) glycerol in the absence or presence of 1 mg/mL “Ceres”. The absorbance signals were normalized using the CH_2_ asymmetric C–H stretch peak of PE (≈2,913 cm^-1^) as the reference (not visible in this zoomed-in view). See Fig. S7 for the complete FTIR dataset used to produce this figure. The data were acquired with 8 cm^-1^ resolution. The same results were obtained when repeating the experiment using a higher (3 mg/mL) concentration of “Ceres” and recording the FTIR data with higher (2 cm^-1^) resolution (Fig. S8).

Next, the molar mass distributions in “Ceres”-treated and buffer-treated LDPE films were analyzed using SEC (Fig. 6). The results clearly show that incubating “Ceres” with LDPE did not result in any deconstruction. The experiments above were conducted using chemically pure LDPE films, produced in-house, whereas Sanluis-Verdes *et al*. used commercial PE film of undescribed origin, likely to contain additives, such as antioxidants, slip agents, or other stabilizers. Sanluis Verdes *et al*. speculated that the degradation of PE by hexamerins from *Galleria mellonella* may depend on the presence of some additives acting as redox mediators. Therefore, we repeated our FTIR experiments using a commercial LDPE film (see Materials for the detailed description) instead of pure LDPE film. No signs of PE oxidation by “Ceres” were observed despite testing multiple reaction conditions (pH 5.0, pH 7.0 and pH 9.0; Fig. S9).

**Figure 6.**
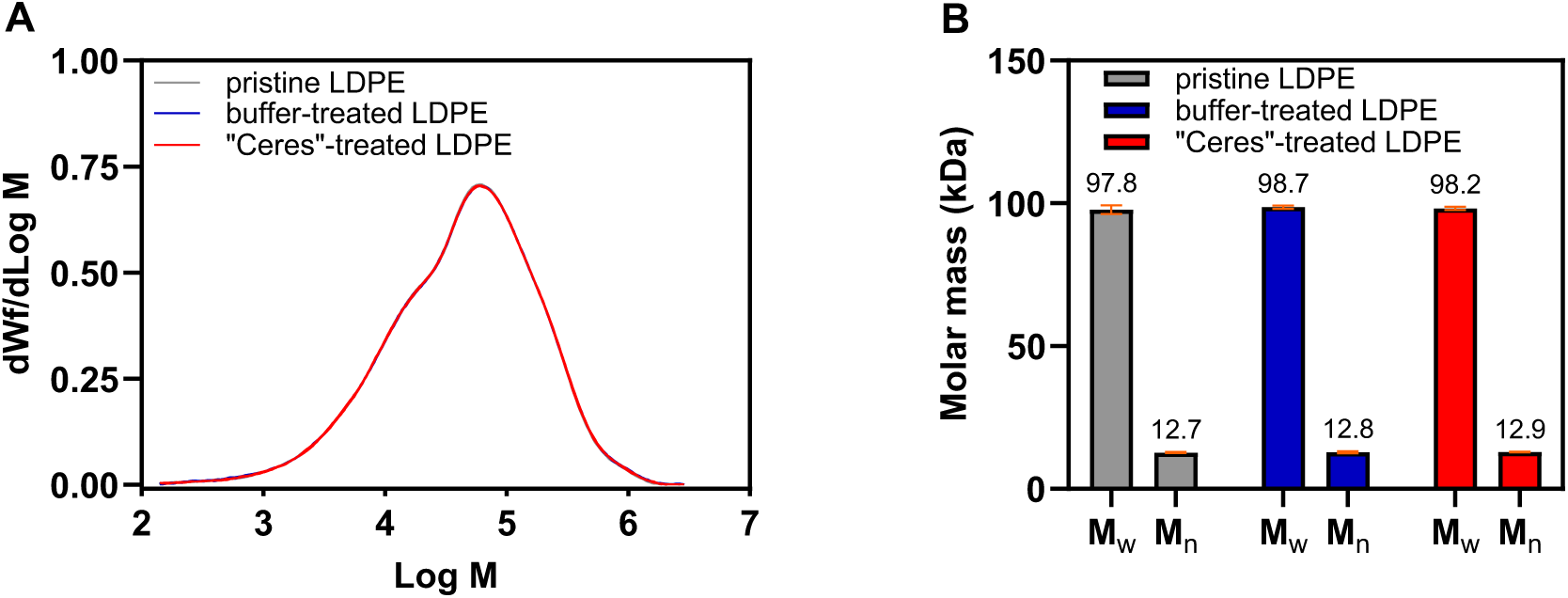
Effect of incubation with “Ceres” on the molar mass distribution in LDPE. The figure shows the results of SEC analysis of LDPE films that had been treated with “Ceres” and of LDPE films from control reactions. To acquire data for each sample type, 9 polymer film discs from 9 independent incubations were pooled together, dissolved in 1,2,4-trichlorobenzene and analyzed three times. Note that panel A features nine molar mass distribution curves (three traces per sample type) that are overlapping almost completely. The error bars in panel B represent standard deviations between technical triplicates. The experiments were conducted for 70 hours at 25 °C in the absence or presence of 1 mg/mL “Ceres” using in-house made additive-free LDPE films fully submerged in 20 mM HEPES, pH 7.0, supplied with 150 mM NaCl and 5% (v/v) glycerol. The pristine LDPE samples were not subjected to any type of treatment. M_w_, weight average molar mass; M_n_, number average molar mass.

In a final attempt to detect “Ceres”-dependent oxidation of polyethylene, the treated commercial LDPE samples were subjected to pyrolysis-gas chromatography/mass spectrometry (Py-GC/MS). This highly sensitive analytical approach has been successfully used to demonstrate abiotic oxidation of PE by UV light [35]. The volatile products released from “Ceres”-treated samples upon thermal degradation were identical to products released from the buffer-treated films (Fig. S10) and not indicative of any enzymatic surface modification of plastic. Taken together, our SEC, Py-GC/MS, and FTIR data suggest that “Ceres” is not capable of oxidation and deconstruction of LDPE films.

### Wax worm saliva displays phenoloxidase activity but does not cause PE deconstruction

Not being able to observe PE oxidation by recombinantly produced “Ceres”, we treated LDPE samples with crude saliva of the greater wax moth larvae (GmSal) that, according to Sanluis-Verdes *et al*. [23], contains both “Ceres” and another plastic-active hexamerin, “Demetra”. Heat inactivation of enzymes in insect salivary samples is a critical control to ensure that observed signs of LDPE film oxidation are due to enzyme activity. To the best of our knowledge, such control experiments have not been reported in studies claiming the presence of PE-degrading enzymes in GmSal. To ensure extracted GmSal has enzymatic activity and to confirm the effectiveness of enzyme heat inactivation, a phenoloxidase (PO) activity assay was performed. The assessment of PO activity was carried out with increasing volumes of GmSal; as GmSal volume increased activity on L-DOPA increased, indicating the presence of active POs in the saliva (Fig. 7). Model PO mushroom tyrosinase from *Agaricus bisporus* was used as a positive control to confirm PO activity. Importantly, when heat inactivated at 95 °C for 15 min, both the positive control tyrosinase and GmSal showed no residual PO activity, confirming the heat inactivation method is effective.

**Figure 7.**
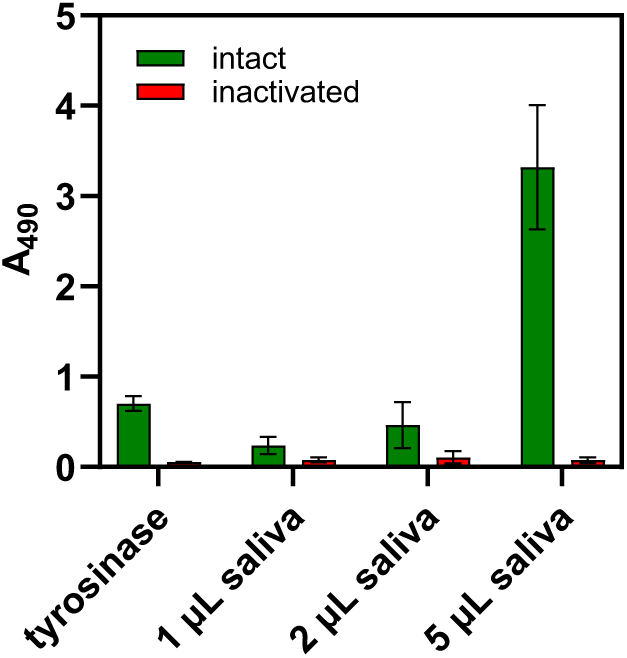
Phenoloxidase activity in wax worm saliva (GmSal) before and after heat inactivation. The graph shows endpoint data from 12-hour reactions and demonstrates the conversion of L-DOPA to dopachrome, which is reflected in an increase in absorbance at 490 nm. As saliva concentration is increased, dopachrome production increases, indicating phenoloxidase activity. Note that the reactions with heat-treated saliva (“inactivated”) show essentially no conversion of L-DOPA. The experiments were carried out at 24 °C using 1 µM of tyrosinase from *Agaricus bisporus* (positive control) or 1-5 µL of GmSal in 20 mM HEPES, pH 7.0, supplied with 4 mM L-DOPA, 150 mM NaCl and 5% v/v glycerol. Error bars indicate standard deviations between triplicates.

The protocol for assessing GmSal-catalyzed changes on the molar mass of plastic substrates presented by Sanluis-Verdes *et al*. was replicated using an additive-free LDPE film. After fifteen consecutive applications of 100 µL of GmSal, with 90 minutes of incubation time after each application, no changes in the molar mass distribution were observed (Fig. 8). Neither intact nor heat-inactivated saliva resulted in a change in the weight-average molar mass of the films post treatment. Importantly, the unprocessed additive-free PE film used in our study falls within the molar mass range of the commercial PE film used in the original study, implying that the original study did not use a PE with shorter chains that could be easier to degrade.

**Figure 8.**
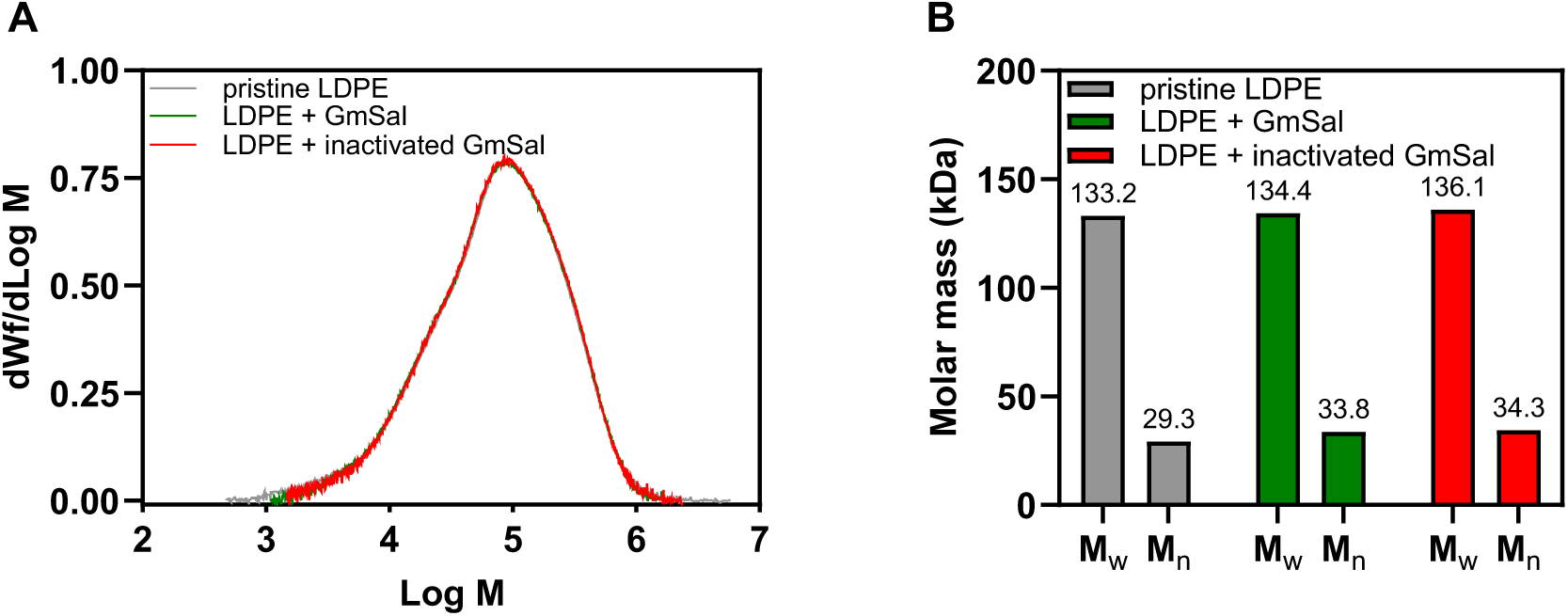
Effect of wax worm saliva (GmSal) on the molar mass distribution in additive-free LDPE film. The figure shows the results of SEC analysis of films treated with the intact or heat-inactivated GmSal and of the pristine material. 100 µL of saliva was applied to the centre of a 5 mm x 5 mm film (i.e., approximately the same position each time) 15 times and the film was incubated for 90 minutes at 24 °C between each application. M_w_, weight average molar mass; M_n_, number average molar mass.

### Revisiting previously reported FTIR data: PE oxidation by wax worm saliva is a false positive

To further assess the PE-degrading potential of wax worm saliva, we again used additive-free LDPE films. The use of additive-free films ensures that observed apparent changes in the plastic are true changes to the chemistry of PE. The GmSal treatments done prior to FTIR measurements mirrored the approach taken by Sanluis-Verdes *et al*., but with additional control experiments that were omitted in the original study. Firstly, we treated LDPE with thermally denatured GmSal. Secondly, we also studied the FTIR spectrum of GmSal alone, since we suspected that the FTIR spectrum could be affected by components in the saliva. After nine consecutive applications of 5 µL of intact or heat-inactivated GmSal (with a protein concentration of approximately 25 mg/mL, i.e., in the concentration range reported in the original study), with 90 minutes incubation at 24 °C after each application, the films were washed with water and 70% (v/v) EtOH and were allowed to air-dry. The FTIR spectrum showed typical features indicative of protein related contaminants being bound to the film. N-H stretching signals at approximately 3,300 cm^-1^ and amide I and II peaks around 1,650 cm^-1^ and 1,550 cm^-1^ were observed for the films treated with both active and heat-inactivated GmSal (Fig. 9). These peaks are characteristic of protein adhered to the film [36] and do not reflect chemical modification of PE. This is confirmed by the FTIR spectrum of GmSal alone, which also shows these peaks (Fig. 9). Additionally, a peak at 1,080 cm^-1^ is visible in spectra for the films treated with GmSal and heat-inactivated GmSal and in the spectrum for GmSal alone. Since this peak appears in all spectra, it must reflect a salivary compound that is bound to the film.

**Figure 9.**
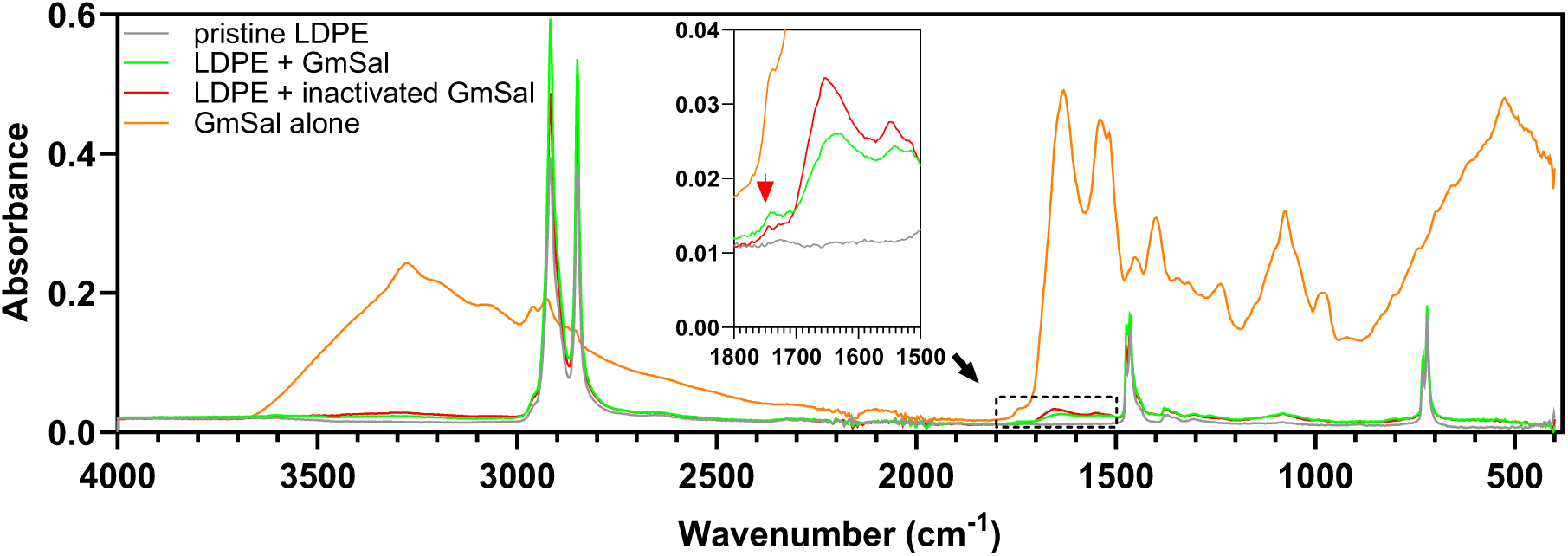
FTIR spectra of additive-free LDPE films subjected to treatment with wax worm saliva (GmSal). The spectra are for an untreated control film (grey), GmSal-treated PE (green), PE treated with heat-inactivated GmSal (red), and GmSal alone (orange). The spectra for treated PE and for GmSal alone show peaks corresponding to protein (N–H stretch around 3,300 cm^-1^, amide I and II peaks at 1,650 and 1,550 cm^-1^), and a peak at approximately 1,080 cm^-1^, likely reflecting a hydrophobic contaminant(s) from the saliva. The peak at ≈1,745 cm^-1^ featured in the zoomed-in view and indicated with a red arrow is also present in all spectra except untreated PE and likely represents a saliva compound; see text for more details. 5 mm x 5 mm PE films were treated by nine consecutive applications of 5 µL of GmSal to approximately the same position in the centre of the film, with 90 minutes incubation at 24 °C between applications. To obtain the spectrum of GmSal alone, the liquid was allowed to air-dry, and the dry matter was deposited to the ATR sensor.

In the original report by Sanluis-Verdes *et al*., a minor peak around 1,745 cm^-1^ that appears upon treating PE film with saliva is highlighted and claimed to demonstrate oxidation of the film. This peak is indeed consistent with carbonyl groups and could thus suggest oxidation, and we do indeed observe this peak after treating our treated LDPE with GmSal. However, importantly, this peak also appears when analyzing GmSal alone (Fig. 9). Thus, this signal likely reflects a contaminating compound in GmSal. Moreover, this peak is present regardless of whether the GmSal was heat-treated or not, making it highly improbable that the signal is related to enzymatic modification of the plastic.

## Discussion

In this work, two recently proposed plastic-active enzymes were produced recombinantly using expression platforms similar to those reported in the original papers by Zhang *et al*. and Sanluis-Verdes *et al*. [22, 23]. While we replicated some of the original observations, our comprehensive analyses, involving multiple reaction conditions and additional control experiments and analytical techniques not featured in the initial studies, show that these enzymes exhibit no activity on PVC or PE, respectively.

Regarding the catalase-peroxidase from *Klebsiella* sp. EMBL-1 (*Kleb*CP), our data clearly indicate that the produced enzyme is active and capable of peroxidation of a model small molecule substrate (Amplex Red) and of H_2_O_2_ turnover via a catalase pathway. At the same time, *Kleb*CP produced in our laboratory is not able to cause deconstruction or surface oxidation of the two types of PVC tested. Importantly, our spectroscopy experiments with *Kleb*CP in the absence of plastics strongly suggest that the FTIR data reported by Zhang *et al*. [22] as the evidence for oxidation of PVC constitutes a false positive, caused by protein adsorption to the polymer surface.

If one is willing to accept the previous claim that *Kleb*CP is capable to cause PVC peroxidation and deconstruction, it is vital to understand how such reactions are possible under the conditions used by Zhang *et al*., which did not involve the essential peroxidase co-substrate H_2_O_2_. Although our data indicate the possible presence of redox active compounds in PVC powders that can act as a weak *in situ* H_2_O_2_ source by reacting with O_2_, the plastic samples featured in the Zhang *et al*. study were purified by using a dilution-precipitation approach to remove any additives. The apparent lack of a proper co-substrate in the proposed PVC degradation reactions was not addressed in the original study, making interpretation of the results difficult. It cannot be excluded that *Kleb*CP is able to use O_2_ for substrate oxidation in the absence of H_2_O_2_, thus acting as oxidase rather than a peroxidase [37]. However, evidence for oxidase activity in peroxidases is scarce.

It is worth noting that the Zhang *et al*. study presented a GC-MS analysis of potential products of PVC degradation by *Kleb*CP. A set of C_20_, C_21_, C_24_ and C_25_ alkanes was detected in dichloromethane extracts of the reaction supernatants. Although our own experiments do not address this observation, it seems unlikely that a single enzyme will both cleave C–C bonds and remove Cl to produce the reported fully dechlorinated products. According to Zhang *et al*., the *Klebsiella* sp. EMBL-1 strain can use PVC as a sole carbon source. We cannot exclude that this bacterium can indeed deconstruct and metabolize PVC using yet to be identified enzymes, but our data clearly show that *Kleb*CP is unlikely to play a central role in this process.

Our experiments with a recombinantly produced hexamerin from *Galleria mellonella* (“Ceres”) revealed no activity on PE. Insect hexamerins are phylogenetically related to phenoloxidases, which has led to the postulation that “Ceres” (alongside another insect hexamerin, “Demetra”) possesses a “phenoloxidase-like” activity, allowing for PE oxidation and depolymerization via an unknown mechanism [23]. It was also proposed that such a mechanism may involve hexamerin-dependent oxidation of small molecule additives in commercial PE, generating redox mediators and promoting radical reactions on the polymer. In our view, there are several facts that make such a line of reasoning implausible. While insect hexamerins and phenoloxidases indeed share structural similarity, hexamerins lack the canonical copper-binding motif (six histidine residues coordinating two copper atoms) characteristic of phenoloxidases and essential for the interaction with oxygen and the oxidase activity [31] (Fig. S3), and neither “Ceres” nor “Demetra” have a phenoloxidase-like catalytic site. This discrepancy between the suggested activity and the sequence of the proteins in question was acknowledged by the authors from the Sanluis-Verdes *et al*. paper in a very recent publication [30]. The structural model of “Ceres” proposed in this recent paper suggests the presence of two potential non-canonical metal binding sites per protein molecule but does not describe the nature of these metal atoms. Our ICP-MS analysis of “Ceres” showed that this hexamerin does not bind copper, meaning that any reference to phenoloxidases while discussing “Ceres” (e.g., using the term “phenoloxidase-like activity”) is potentially misleading. Not surprisingly, “Ceres” is not able to act on L-DOPA, a common natural phenoloxidase substrate.

On another note, the radical-based mechanism behind the catalytic action of “Ceres” suggested by Sanluis-Verdes *et al*. [23] does not seem to be compatible with FTIR data presented in their paper. Radical reactions typically lead to formation of a wide range of products, as illustrated by the well-studied processes of PE auto-oxidation and photo-oxidation [38, 39]. For example, FTIR spectra of photo-oxidized PE samples feature broad carbonyl peaks, which can be deconvoluted to reveal the presence of conjugated ketones (≈1,695 cm^-1^), carboxylic acids (≈1,713 cm^-1^), ketones (≈1,720 cm^-1^), esters (≈1,735 cm^-1^), peracids (≈1,765 cm^-1^) and lactones (≈1,785 cm^-1^) [39]. At the same time, PE oxidation observed by Sanluis-Verdes *et al*. after treating PE films with “Ceres” resulted in a narrow signal centred around ≈1,745 cm^-1^, which is indicative of carbonyl groups in esters. It seems unlikely that a radical reaction on PE can lead to the formation of a single type of product. It is also unlikely that concomitant introduction of carbonyl functionalities and C–O bonds into the PE chain, i.e. introduction of ester bonds, can be achieved via a single-enzyme reaction using “Ceres” or any other type of PE-active protein. Finally, the suggestion by Sanluis-Verdes *et al*. that some PE additives can promote radical reactions in the presence of “Ceres” is counterintuitive, given that such compounds are typically used as stabilizers (i.e., it is unlikely that commercial PE additives have strong pro-oxidant properties).

It should be mentioned that the experimental evidence provided by Sanluis-Verdes *et al*. in support of PE oxidation by recombinant versions of proteins found in wax worm saliva is stronger for the other of the two studied hexamerins, “Demetra”. In contrast to “Ceres”, treating PE films with “Demetra” resulted in changes in the surface morphology of PE films. Likewise, putative products of PE deconstruction (2-ketones) were detected by GC-MS only in reactions with “Demetra”, and not in reactions with “Ceres”. Our data clearly show that “Ceres” cannot oxidize or degrade PE and, considering the above, we find it unlikely that “Demetra” can. It is noteworthy that the prevailing view on the functional roles of hexamerins in insects does not entail any enzymatic activity. Hexamerins are thought to be non-catalytic proteins involved in immune responses, binding and transport of organic compounds, and hormonal regulation [40–42].

Interest in “Ceres” and “Demetra” was sparked by the observation that wax worm saliva can degrade PE and that this saliva is rich in these two proteins [23]. Therefore, in parallel with experiments with recombinantly produced “Ceres”, we tried to reproduce this underlying key observation. Our SEC data do not provide support for PE deconstruction by saliva. Furthermore, our additional control experiments using saliva samples lacking any plastics clearly indicate that the ≈1,745 cm^-1^ FTIR spectroscopy peak that has been used by Sanluis-Verdes *et al*. in support of PE oxidation originates from a contaminating compound and not from a true product. This signal could for example reflect the presence of lipids [43, 44]. While it is not clear whether the wax moth larvae used by Sanluis-Verdes *et al*. were fed beeswax at any point during cultivation, this feeding technique is sometimes used according to literature [45]. Beeswax is known to display an FTIR signal around 1,740 cm^-1^ due to the presence of esters [28]. The idea that the apparent plastic deconstruction by wax moth larvae may represent an artefact is not new and was first proposed by Weber *et al*. [46]. While our study does not address the capacity of this insect to use PE as a carbon source, it is important to note that direct evidence for plastic metabolization by *Galleria mellonella* (e.g., tracing experiments showing incorporation of isotopes from labelled PE into larval tissues) has not yet been reported to our knowledge.

We cannot fully rule out that plastic deconstruction by *Kleb*CP, “Ceres”, and/or crude saliva of *Galleria mellonella* larvae may happen, as it may depend on substrate properties, such as the degree of crystallinity, form factor, or the presence (or absence) of small molecule additives. For example, in contrast to our work, the PVC samples used by Zhang *et al*. to test the activity of *Kleb*CP were produced by precipitating the polymer from a tetrahydrofuran solution, potentially resulting in small and refined particles with a very high surface available to an enzyme. On another note, the FTIR spectrum of the commercial PE film used by Sanluis-Verdes *et al*. features a few strong additional peaks (≈875 cm^-1^, ≈1,425 cm^-1^; see the source data file provided for Fig. S8 in the original manuscript [23]), indicating the presence of some additive(s) that may affect substrate properties. The lack of detailed information on the model plastics (including information on the origin of the materials and their characterization) in the studies conducted by Zhang *et al*. and Sanluis-Verdes *et al*. complicates further analysis. In the most optimistic scenario, if one chooses to accept the original claims that *Kleb*CP and “Ceres” can oxidize plastics, the lack of any PE and PVC deconstruction observed in our own study under multiple reaction conditions with several different substrates should be taken as a sign that these enzymes are not robust and/or only act on a limited subset of possible PVC/PE substrates, which casts doubt on their near-term practical application.

Establishing efficient enzymatic conversion of non-hydrolyzable polymers such as PVC and PE would be a major achievement, and it is no wonder that signs of such conversion and the enzymes potentially involved are receiving considerable attention. The idea that non-hydrolysable polymers can be degraded using enzymes was put forward long ago. For example, in 2001 Fujisawa *et al*. reported a remarkable 8-fold decrease in the (weight average) molar mass of a non-pretreated PE membrane, after just 3 days of incubation with *T. versicolor* laccase and a redox mediator (HBT) [12]. A recent review on PE biodegradation by Jin *et al*. [47] refers to 12 proteins potentially capable of enzymatic activity on PE (2 of which are “Ceres” and “Demetra”). Yet, despite previous claims, enzymatic degradation of PE remains controversial and our present data show that the conclusions drawn from recent and seemingly groundbreaking work with PE and PVC [22, 23] were too optimistic. Clearly, the field would benefit from standardization and rigorous validation of analytical methods including absolute requirements for control reactions. With much research ongoing, time will tell if Nature manages effective biological deconstruction of non-hydrolyzable plastics using catalytic tools that are amenable to industrial use.

## Methods

### Materials

Chemicals were obtained from Sigma-Aldrich (St. Louis, MO, USA) unless specified otherwise. Amplex Red (10-acetyl-3,7-dihydroxyphenoxazine) was purchased from Thermo Fisher Scientific (Waltham, MA, USA), dissolved in DMSO at 10 mM final concentration and stored at -20 °C as 50 µL aliquots in PCR tubes, protected from light. Mushroom tyrosinase was supplied by Sigma-Aldrich (catalog number T3824-25KU). L-DOPA was obtained from Cayman Chemical Company (Ann Arbor, MI, USA) or from Sigma-Aldrich (St. Louis, MO, USA). Butylated hydroxytoluene was purchased from Alfa Aesar (Haverhill, MA, USA) at 99% purity. 1,2,4-trichlorobenzene (HPLC grade) was purchased from Thermo Fisher Scientific. Tetrahydrofuran (HPLC grade, stabilized with BHT) was purchased from Spectrum Chemical (New Brunswick, NJ, USA).

Two types of PVC powder were purchased from Sigma-Aldrich to be used as model catalase-peroxidase substrates: “low molecular weight PVC” and “high molecular weight PVC” (catalog numbers 81388 and 81387, respectively). These samples were shown to be fully amorphous, according to differential scanning calorimetry (DSC; see below). The molar mass distribution in both PVC samples were determined by size exclusion chromatography (see below for the method description and see Fig. 2 for the SEC data).

Two types of PE films were used to assess the activity of “Ceres”:

1. Additive-free 0.05 mm LDPE (low density polyethylene) film produced in-house by blown film extrusion process from Borealis FT5230 pure LDPE resin (Borealis, Vienna, Austria) using a laboratory scale film extruder E25P x 25 L/D (Collin Lab & Pilot Solutions GmbH, Maitenbeth, Germany) at a processing temperature of 175 °C. The crystallinity of this film was determined to be 44.7 ± 0.1%, using DSC.
2. Commercial transparent LDPE film obtained from external packaging supplied with 50 mL polypropylene centrifuge tubes (Grenier, Kremsmünster, Austria; Sigma-Aldrich catalog number T2318). The crystallinity of the film was determined to be 43.7 ± 0.1%, using DSC.

Additive-free 0.035 mm LDPE film obtained from GoodFellow Cambridge Limited (Huntington, UK) was used to study PE-degrading activity in the saliva of *Galleria mellonella* larvae (waxworms). The crystallinity of the film was determined to be 41.7 ± 3.3%, using DSC. Waxworms were obtained from Rainbow Mealworms (Compton, CA, USA).

### Differential scanning calorimetry of polymer samples

The crystallinity of plastic samples was assessed using DSC. Most of the materials (except LDPE films involved in Fig. 8,9) were analyzed with a DSC250 calorimeter (TA instruments, New Castle, DE, USA). About 3-5 mg of the samples was encapsulated into sealed aluminium pans and subjected to heating from -10 °C to 200 °C at a heating rate of 10 °C/min followed by an isothermal hold of 5 min. The percentage of crystallinity in the polymer resins was determined according to the following equation:

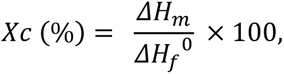

where *ΔH_m_* is the enthalpy of melting derived from the thermogram and 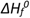 is a reference value for the enthalpy of melting of 100% crystalline material (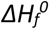 is 293 J g^-1^ for PE) For each sample, the analysis was carried out in duplicates and average values are reported.

LDPE films involved in the experiments shown in Fig. 8, 9 were analyzed under a N_2_ environment with a Discovery DSC machine (TA Instruments, New Castle, DE) by heating the sample at 20 °C/min from 25 °C to 250 °C. The resulting thermograms were used for crystallinity calculations according to the equation above.

### Size exclusion chromatography (SEC) of polymer samples

The weight average molar mass (M_w_), number average molar mass (M_n_) and the molar mass distribution in PVC materials were determined using a 1260 HPLC system equipped with three PLgel 5 µm Mixed-C columns and a PLgel guard column (Agilent Technologies, Santa Clara, CA, USA). Approximately 5 mg of PVC was dissolved in 1 mL of tetrahydrofuran at room temperature for at least 14 h. Samples were 0.2 µm filtered, and 100 µL was injected into the SEC system. The analysis was performed at 40 °C using an Optilab differential refractive index detector and a miniDAWN multi-angle light scattering detector (Wyatt Technology, Santa Barbara, CA, USA) with tetrahydrofuran stabilized with BHT as the mobile phase (1 mL/min flow rate). A polystyrene standard with a narrow molecular mass distribution (Wyatt Technology) was used for detector calibration and to confirm system performance. Data analysis was performed in Astra (Wyatt Technology) with a dn/dc value for PVC in THF of 0.0961 [48].

The weight average molar mass (M_w_), number average molar mass (M_n_) and the molar mass distribution in PE materials (except the LDPE films involved in Fig. 8,9) were determined using a GPC-IR5 system (Polymer Char, Valencia, Spain) equipped with four PLgel 20 µm MIXED-A columns (Agilent Technologies, Santa Clara, CA, USA). Approximately 4 mg of LDPE film was pre-dissolved in 8 mL of 1,2,4-trichlorobenzene at 160 °C for 3 hours and 200 µL samples were injected into the SEC system. The analysis was performed at 150 °C using a high sensitivity infrared detector and 1,2,4-trichlorobenzene as a mobile phase (1 mL/min flow rate). Polystyrene standards with a narrow molecular mass distribution and M_peak_ in the range of 1,140 – 7,500,000 g/mol (Agilent Technologies, Santa Clara, CA, USA) were used for calibration.

The SEC analysis of the samples used to produce Fig. 8 and Fig. 9 was carried out on TOSOH HLC-8312GPC/H system equipped with two TSKgel GMHHR-H(20)HT columns and one TSKgel G2000HHR column in series, coupled with refractive index (RI) and viscosity detectors (TOSOH Biosciences, PA, USA). HPLC-grade 1,2,4-trichlorobenzene stabilized with 500 ppm butylated hydroxytoluene was used as the mobile phase. LDPE samples (3–10 mg) were dissolved in mobile phase (3–6 mL) at 140 °C for a minimum of two hours to generate nominal concentrations of 2 mg/mL. 300 µL of samples were injected and eluted for 80 minutes at a flow rate of 0.8 mL/min at 140 °C. Molar mass distributions were determined using the RI responses, and SEC samples were calibrated against linear polystyrene standards (9 runs between 6×10^2^–2×10^6^ g/mol). Molar masses of LDPE obtained were corrected using the Mark-Houwink relationship (following previously detailed methods) [49].

### FTIR spectroscopy analysis of polymer samples

All FTIR spectroscopy data in this study, except spectra shown in Fig. 9 were collected with a 5500 Series spectrometer equipped with a diamond-ATR (attenuated total reflectance) sampling module (Agilent Technologies, Santa Clara, CA, USA). The signals were obtained with 8 cm^-1^ spectral resolution (unless specified otherwise) by performing 35 consecutive readings per measurement in 4,000-650 cm^-1^ range using MicroLab PC software (Agilent Technologies) and further processed with Spectragryph (F. Menges, “Spectragryph - optical spectroscopy software”, Version 1.2.16, 2023, http://www.effemm2.de/spectragryph/) to perform the automatic baseline correction and signal normalization (see Figure legends for more details).

The data featured in Fig. 9 were acquired with a Thermo Scientific Nicolet iS5 FTIR spectrometer equipped with a diamond-ATR cell (Pittsburgh, PA, USA). Spectra were recorded in the range of wavelengths 4,000−400 cm^−1^ with a minimum of 32 scans per measurement and a spectral resolution of 0.482 cm^−1^.

### Cloning, expression and purification of *Kleb*CP

To retrieve the gene encoding for *Kleb*CP, the cloning primers reported by Zhang *et al*. [22] (F: 5′-ACGAATTCATGAGCACGTCTAACGAC-3′, R: 5′-AACTCGAGCAGGTCGAAGCGGTCGA-3′) were searched against the complete genome of *Klebsiella* sp. strain EMBL-1 (NCBI accession number CP079802.1) using PrimerBLAST [50]. The protein sequence of catalase-peroxidase was derived from the identified gene (NCBI accession number QXX04157.1). The DNA sequence encoding for the enzyme with a C-terminal purification tag (“AHHHHHH”) was codon-optimized for expression in *Escherichia coli*, synthesized and cloned into the pET-26b(+) expression vector using NdeI/XhoI restriction sites by GenScript (Piscataway, NJ, USA). The resulting vector was used to transform One Shot™ BL21 Star™ (DE3) chemically competent *E. coli* cells according to manufacturer’s protocol. Transformants were selected by growing on LB agar plates containing 50 µg/mL kanamycin, overnight. A single colony was then picked and grown in 10 mL of LB medium containing 50 µg/mL kanamycin at 37 °C and 200 rpm overnight. 5 mL of this pre-culture were used to inoculate 1 L of LB medium containing the same concentration of kanamycin, which was incubated at 37 °C and 200 rpm until reaching an OD_600_ of approximately 0.6. Next, expression of *Kleb*CP was induced by adding isopropyl thiogalactopyranoside (IPTG) to a final concentration of 0.5 mM and the culture was further incubated at 30 °C and 200 rpm for 16 h. The cells were harvested by centrifugation at 6,000 g for 15 minutes and re-suspended in 30 mL binding buffer (500 mM NaCl, 10 mM imidazole, 20 mM Tris, pH 8.0). The suspension was sonicated using a VibraCell ultrasonic disintegrator equipped with a micro tip probe (Sonics, Newtown, CT, USA). The amplitude was set to 29% and the sonication was performed for 10 minutes in 5 second steps followed by 5 second pauses. The lysate was clarified by centrifugation at 20000 g for 15 minutes at 4 °C and filtered through a 0.22 µm syringe filter. The enzyme was purified using a 5-mL Ni-charged EconoFit Nuvia IMAC Column (Bio-Rad, Hercules, CA, USA) and a linear gradient of 10 - 250 mM imidazole in binding buffer (100 mL in total at 2.5 mL/min flow rate). The resulting enzyme preparation was analyzed by SDS-PAGE and the concentration of the enzyme was estimated by measuring the absorbance at 280 nm using the theoretical extinction coefficient of *KlebCP* (143,475 M^-1^ cm^-1^) calculated with ProtParam [51].

### Peroxidase activity of *KlebCP* on Amplex Red

*Kleb*CP was tested for peroxidase activity using the Amplex Red reagent (Thermo Fisher Scientific, Waltham, MA, USA) by following the formation of a highly fluorescent product, resorufin. The assay was carried out in a 96-well plate using a VarioSkan Lux microplate reader (Thermo Fisher Scientific). The reaction mixture consisted of 300 mM NaCl, 0.1 μM *Kleb*CP and 100 µM Amplex Red in 50 mM Tris-HCl buffer, pH 8.0, with a final volume of 100 µL. The reaction was initiated by adding H_2_O_2_ to a final concentration of 100 µM, and the fluorescence (excitation 560 nm, emission 585 nm) was measured every 5 s for 2 minutes while incubating at 30 °C in the microplate reader. Within this time, a plateau was reached, and further H_2_O_2_ addition steps were performed to restart the reaction, as shown in Fig. 1. In total, H_2_O_2_ was introduced to the reaction mixtures 3 times.

### Treatment of PVC samples with *Kleb*CP

The activity of *Kleb*CP on plastics was tested using two types of PVC powder available via Sigma-Aldrich (St. Louis, MO, USA): “low molecular weight PVC” and “high molecular weight PVC” (catalog numbers 81388 and 81387, respectively). The reactions were carried out in a final volume of 1000 µL, containing 50 mM Tris-HCl buffer, pH 8.0, 300 mM NaCl, 100 mg/mL PVC powder and 10 µM *Kleb*CP. Reaction mixtures lacking *Kleb*CP were used as negative controls. The tubes were incubated in an Ecotron shaker (Infors HT, Bottmingen, Switzerland) at 30 °C and 150 rpm for 96 h. The liquid phase was removed by pipetting, and the PVC particles were dried at 80 °C for 2 h and subjected to FTIR spectroscopy, as described above. Additional washing step was performed to remove residual *Kleb*CP, adsorbed on PVC surface, by adding 1000 µL Milli-Q water and incubating for 30 min at 30 °C and 1000 rpm in a ThermoMixer (Eppendorf, Hamburg, Germany). Then, the liquid phase was removed by pipetting and the PVC particles were dried for 2 h at 80 °C. Another set of FTIR spectra was acquired after washing.

To obtain the FTIR spectrum of *Kleb*CP alone, 6 µL of a 66 mg/mL enzyme solution was mixed with 500 µL of absolute ethanol to precipitate the protein. Ten 2 µL drops of the suspension were sequentially applied to the surface of an ATR-sensor and allowed to dry before recording the signal.

To produce PVC samples for SEC analysis, reactions with plastic powders and *Kleb*CP were set up as described above. Additional reactions were set up that contained 50 nM of an engineered cellobiose dehydrogenase (CDH) [26] and 5 mM cellobiose, to provide an *in situ* H_2_O*_2_* source for the catalase-peroxidase. After treatment, the PVC material was washed with 1 mL Milli-Q water and dried, as described above.

### Detection of H_2_O_2_ formation under reaction conditions used in PVC degradation experiments

The accumulation of H_2_O_2_ in reaction mixtures was assessed using the HRP/Amplex Red assay [52]. Reaction mixtures containing 50 mM Tris-HCl buffer pH 8.0 and 300 mM NaCl were supplied with 100 µM Amplex Red and 5 U/mL horseradish peroxidase (HRP). These experiments were conducted in the presence of 100 mg/mL PVC particles and in the presence or absence of 50 nM CDH and 5 mM cellobiose. The apparent generation of H_2_O_2_ was detected at room temperature by following the formation of resorufin (excitation 560 nm, emission 585 nm) using a VarioSkan Lux microplate reader (Thermo Fisher Scientific, Waltham, MA, USA).

### Cloning, expression and purification of “Ceres”

Cloning, production, and purification of “Ceres” were performed by GenScript (Piscataway, NJ, USA). In brief, the DNA sequence corresponding to the “Ceres” protein (NCBI accession number XP_026756459.1) was codon optimized for expression in an Sf9 (*Spodoptera frugiperda*) insect cell line. This sequence was further modified by substituting the native signal peptide with the gp64 secretion signal of baculoviral origin (“MLLVNQSHQGFNKEHTSKMVSAIVLYVLLAAAAHSAFA” [53]) and by incorporating a short downstream element encoding for C-terminal affinity tag (“AHHHHHH”), followed by a stop codon. The resulting DNA was synthesized and cloned into the pFastBac1 vector using EcoRI/HindIII restriction sites. This donor plasmid was then used to transform DH10Bac *E. coli* cells containing a baculovirus shuttle vector to produce a “Ceres”-encoding bacmid through recombination. The bacmid was purified from the *E. coli* culture and used to transfect Sf9 cells, leading to the formation and release of viral particles carrying the target gene. To produce “Ceres”, the obtained viral particles were used to infect a fresh Sf9 cell culture that was incubated in 2 L of Sf-900 II SFM medium for 3 days at 27 °C. The secreted mature protein was purified from clarified culture medium using metal affinity chromatography, shipped in a dry ice in 50 mM Tris-HCl, pH 8.0, and stored at -80 °C until further use.

### LC-MS/MS analysis of the purified “Ceres” protein

Approximately 0.5 µg of purified “Ceres” protein was subjected to SDS-PAGE. The gel was stained with Coomassie R-250 (Imperial Protein Stain, Thermo Fisher Scientific, Waltham, MA, USA) and de-stained according to the manufacturer’s instructions. Next, the “Ceres” band (≈ 80 kDa) was cut out with a scalpel and incubated with 10 mM dithiotreitol for 30 min at 56 °C, followed by alkylation with 50 mM iodoacetamide for 30 min, at room temperature, in the dark. Finally, in-gel digestion using 10 ng/µL of sequencing grade modified trypsin (Promega, Madison, WI, USA) was performed overnight at 37 °C.

The peptides were extracted from the gel by acidifying the solution with trifluoroacetic acid (TFA) and sonicating for 10 min in an ultrasonic water bath 3510E-DTH (Branson, Brookfield, USA). The peptides were purified using solid phase extraction pipette tips as described previously [54, 55]. The peptides were then dried in a SpeedVac vacuum concentrator (Thermo Fisher Scientific) and re-dissolved in 0.05% TFA and 2% acetonitrile (ACN). 0.2 µg of peptide sample was analyzed with a RSLCnano UHPLC system coupled to a Q-Exactive mass spectrometer (Thermo Fisher Scientific) using an acetonitrile gradient (3.2% - 60%) in 0.1% formic acid. The mass-spectrometry data were analyzed with the Mascot software (http://www.matrixscience.com), by matching the results to a database [56] containing the full proteome of *Spodoptera frugiperda* (i.e., the expression host) and to the sequence of the mature “Ceres” protein including the purification affinity tag.

### Total copper quantification using inductively coupled plasma mass spectrometry (ICP-MS)

Approximately 450 µL of a 1 µM “Ceres” solution in TraceSELECT metal-free water (Honeywell, Charlotte, NC, USA) was transferred to 20 mL polytetrafluoroethylene vessels. The mass of each “Ceres” sample was measured using an analytical balance (Sartorius, Krugersdorp, South Africa) and converted back to volume by assuming a density of 1 g/mL. Next, 1 mL ultrapure concentrated nitric acid was added using a 5 mL bottle-top dispenser (Seastar Chemicals, Sidney, BC, Canada). Ultrapure nitric acid was produced in-house from trace analysis grade nitric acid (Merck, Darmstadt, Germany) using a SubPur quartz sub-boiling distillation system (Milestone, Italy). The samples were digested using an UltraClave high-performance microwave reactor (Milestone, Italy) by increasing the temperature from 50 °C to 245 °C over 30 min, and then holding 245 °C for 20 min. The digested samples were transferred to 15 mL metal-free polypropylene vials (VWR, Randor, PA, USA) and diluted with ultrapure water to achieve 0.6 M final acid concentration. The weight of the diluted samples was measured with an analytical balance and converted to the final volume (the density of 0.6 M HNO_3_ is 1.0167 g/mL). Total iron and copper concentrations were assessed by a tandem quadrupole 8800 ICP-QQQ system (Agilent Technologies, Santa Clara, CA, USA) using mixed hydrogen and helium as reaction cell gases. Multi-element stock solutions (LabKings B.V., Hilversum, The Netherlands) were used for the instrument calibration and quality control. An internal standard containing 10 μg/L of Rh was automatically mixed with the sample in the prepFAST system (Elemental Scientific, Omaha, NE, USA). Two independently prepared “Ceres” samples were analyzed with each sample injected twice.

### Assessing enzymatic oxidation of L-DOPA

The capacity of “Ceres” to catalyze the oxidation of L-3,4-dihydroxyphenylalanine (L-DOPA) was tested by following the formation of dopachrome. The reactions were carried out in 96-well microtiter plates at 25 °C. The experiments were initiated by adding 10 µL of a “Ceres” stock solution (or 10 µL of Milli-Q water in case of negative control samples) to 190 µL of a 4 mM L-DOPA solution in 20 mM HEPES, pH 7.0, supplied with 150 mM NaCl and 5% (v/v) glycerol. The final concentrations of “Ceres” and L-DOPA amounted to 1 µM and 3.8 mM, respectively. Additional reactions with 1 µM tyrosinase from *Agaricus bisporus* instead of “Ceres” were set up in the same manner to generate a positive control. The solutions were mixed for 10 s at 600 rpm in a Varioscan LUX plate reader (Thermo Fisher Scientific, Waltham, MA, USA). Dopachrome generation was detected by recording the optical absorbance at 490 nm for a period of 72 min.

As another method for assessing oxidative activity on L-DOPA, the consumption of molecular oxygen in reaction mixtures was monitored over time. These experiments were performed in glass vials (‘Reacti-Vial’; Thermo Fisher Scientific; catalog number TS-13221) with air-tight caps equipped with self-sealing silicone septa. The vials containing magnetic stirrer bars were filled with 1234 μL of 20 mM HEPES, pH 7.0, supplied with 4 mM L-DOPA, 150 mM NaCl and 5% (v/v) glycerol, and carefully sealed with the caps. After incubating and stirring the reaction mixtures for approximately 15 minutes (25 °C, 200 RPM), “Ceres” or *Agaricus bisporus* tyrosinase was introduced by injecting 16 µL of stock solution through the septum using a microcapillary syringe (Hamilton, Reno, NV, USA). Protein stock solutions were substituted with reaction buffer to generate negative control experiments. Note that the final reaction volume was chosen to result in a negligible head space in the sealed vessels. The final concentration of L-DOPA amounted to 3.9 mM, whereas the final concentration of both proteins was 0.1 µM. Molecular oxygen levels were measured over time using an OXY-4 oxygen meter equipped with an NTH-PSt1 optical sensor (PreSens, Regensburg, Germany). The microsensor was pre-calibrated by the manufacturer and mounted inside a stainless-steel needle. It was introduced into the reaction vials by piercing the self-sealing septa.

### Treatment of LDPE films with “Ceres”

To produce standardized samples for degradation assays, both in-house made and commercial LDPE films were perforated with an office hole punch, producing regularly shaped plastic discs. The average weight of these discs amounted to 1.7 ± 0.3 mg (*n* = 5) and 1.9 ± 0.5 mg (*n* = 5) for in-house made and commercial films, respectively. The LDPE discs were transferred into 96-well plates (1 film disc per well) using a syringe needle, followed by careful deposition of 100 µL of reaction buffer containing or lacking 1 mg/mL “Ceres” on top of the film. Four different reaction buffers were screened in this study, as indicated in the corresponding figure legends. Note that in this set up, the LDPE discs remained at the bottom of the wells and fully submerged in reaction mixtures despite the density of PE being lower than the density of water. The microtiter plates were closed with a transparent sealing tape to prevent evaporation of buffer solutions and incubated at 25 °C for 70 hours using a thermomixer (Eppendorf, Hamburg, Germany). After the incubation, the discs were removed from the microtiter plate using clean syringe needles. Stainless-steel staples were attached to each disc (1 staple per disc) to increase the weight and simplify the handling of film samples. Next, the film discs were transferred to 2 mL microcentrifuge tubes (1 tube per 3 discs representing triplicates of the same experiment). The films were then incubated for 30 minutes in 1 mL Milli-Q water at room temperature using a thermomixer set to 1000 rpm. The liquid phase was then substituted with another 1 mL of Milli-Q water. The washing procedure was repeated three times in total. Finally, 1 mL EtOH was used instead of water. The samples were dried in 37 °C incubator for approximately 1 hour and subjected to FTIR spectroscopy as described above. Both sides of LDPE discs were analyzed showing no major difference between measurements, hence the two recorded spectra were averaged.

In some cases, an additional washing procedure was carried out prior to FTIR analysis. The film discs in 2 mL microcentrifuge tubes, which had been washed with water and ethanol as described above, were incubated with 1 mL 1 M NaOH at 65 °C, 1000 RPM, for 30 minutes. The samples were then washed three times with 1 mL Milli-Q water and, finally, with 1 mL of EtOH by vortexing, followed by drying in 37 °C incubator for approximately 1 hour.

After performing FTIR spectroscopy, a sub-set of film samples was subjected to Py-GC/MS analysis, as described below.

For the assessment of the “Ceres”-induced changes in the molar mass distribution in PE, nine independent incubation experiments were carried out in microtiter plates as described above with nine LDPE film discs fully submerged in the reaction buffer containing 1 mg/mL “Ceres”. The films were washed with water and ethanol, and dried as described above, pooled together and dissolved in 1,2,4-trichlorobenzene. The resulting solutions were analyzed with SEC three times (technical triplicates). The same procedure was performed with 9 buffer-treated film discs and 9 pristine film discs, to produce control samples.

As an alternative method to treat LDPE samples with “Ceres”, 5 μL drops of 1 mg/mL protein in 20 mM HEPES, pH 7.0, supplied with 150 mM NaCl and 5% (v/v) glycerol were applied to a 2.5 x 2 cm of the in-house produced LDPE film. The experiment was carried out for 3 days at room temperature with the sample exposed to ambient light. 6 drops of “Ceres” solution were applied to the film with 1 hour interval on each of the first two days, followed by two more drops on the final day (14 x 5 μL solution in total). A stainless-steel staple was attached to the film piece to mark the side that was treated with “Ceres”. The film was washed with 20 mL Milli-Q water (30 minutes) and 20 mL of 50% EtOH (30 minutes) using a rotary shaker (Multi RS-60, BioSan, Riga, Latvia) set to 25 RPM, and then dried for 30 minutes at 37 °C.

### Pyrolysis-gas chromatography/mass spectrometry (Py-GC/MS) analysis of LDPE films

150-250 µg fragments of LDPE film discs were placed in deactivated stainless-steel sample cups and introduced into a multi-shot EGA/PY-3030D pyrolyzer (Frontier Laboratories, Fukushima Japan) coupled to a 7890N gas chromatograph and a 5975 mass selective detector (Agilent Technologies, Santa Clara, CA, USA). The furnace temperature was set to 550 °C and the temperature of the interface between the furnace and the GC/MS system was set to 200 °C. GC injector was operating in split mode (100:1 ratio) at 300 °C. The analysis of volatile products was performed using an Ultra-Alloy metal capillary column containing 5% diphenyl- and 95% dimethylpolysiloxane stationary phase (30 m, 0.25 mm ID, 0.25 µm; Frontier Laboratories Ltd., Koriyama, Japan) using helium as carrier gas at 1 mL/min flow rate. The GC oven temperature was increased from 70 °C to 350 °C at 20 °C/min and maintained at 350 °C for 36 min. The electron ionisation (70 eV) mass selective detector was operating in positive mode with ion source temperature of 230 °C, interface temperature of 300 °C and a scan range of 29-350 m/z.

### Collection of wax worm saliva (GmSal)

*Galleria mellonella* larvae (waxworms) were used for saliva collection. Waxworms were oriented on their dorsal side, with the mouth facing upwards. A 0.5 mm diameter hypodermic needle on the end of a syringe was inserted into the buccal opening of the insect and liquid was drawn outwards using the syringe. Saliva was extracted from each mealworm individually, yielding approximately five to ten microliters per insect. The total protein concentration of pooled saliva samples was measured to be approximately 20 mg/mL using the Bradford assay kit (Bio-Rad, Hercules, CA, USA) and bovine serum albumin (BSA) as standard.

### Assessing the effects of wax worm saliva on L-DOPA and LDPE

To assess activity on L-DOPA and to confirm that heat treatment abolishes this activity, L-DOPA was reacted with saliva with the following reaction conditions: 4 mM L-DOPA, 1-5 µL waxworm saliva, 20 mM HEPES, pH 7.0, 150 mM NaCl, and 5% v/v glycerol. These conditions are the same as those used by Sanluis-Verdes *et al*. in GmSal enzyme studies. 1 µM mushroom tyrosinase (*Agaricus bisporus*) was used as a positive control. Absorbance was measured at 490 nm to observe the formation of dopachrome over the course of 12 hours at 24°C.

Activity of GmSal on PE films was assessed by emulating the approach taken by Sanluis-Verdes *et al.* For FTIR analyses, 5 mm x 5 mm LDPE films (GoodFellow Cambridge Limited, Huntington, UK) were treated by nine consecutive applications of 5 µL of GmSal to the centre of the film (2.5 mm from the edges), with 90 minutes incubation at 24 °C in between applications. For SEC analyses, 100 µL of GmSal was applied to the 5 mm x 5 mm LDPE film 15 times with incubation at 24 °C for 90 minutes in between each application. In both cases, saliva was removed from the film after each incubation period. Prior to FTIR or SEC analyses, each film was washed by vortexing in a 1.5 mL tube for five minutes with 500 µL water and then for five minutes with 500 µL of 70% ethanol. Ethanol was removed and films were allowed to air dry.

## Acknowledgements

We thank Dr. Roland Ludwig at the Department of Food Sciences and Technology, Institute of Biotechnology, University of Natural Resources and Life Sciences (BOKU), Vienna for kindly providing engineered cellobiose dehydrogenase from *Crassicarpon hotsonii* (CDH_oxy+_).

We thank Anica Simic and Kyyas Seyitmuhammedov at the Department of Chemistry, Norwegian University of Science and Technology for ICP-MS analysis of protein samples.

