## Supplementary Figures for "Revisiting the activity of two poly(vinyl chloride)- and polyethylene-degrading enzymes"

### Supplementary information

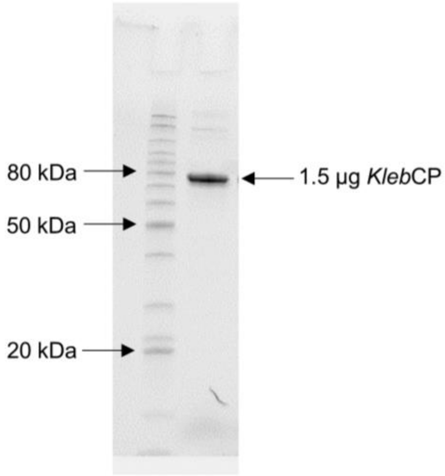


**Figure S1. SDS-PAGE of *Kleb*CP after purification.** The predicted molecular weight of the mature protein with the affinity tag is 79.8 kDa. The molecular weight calculation was carried out using ProtParam tool (https://web.expasy.org/protparam/).

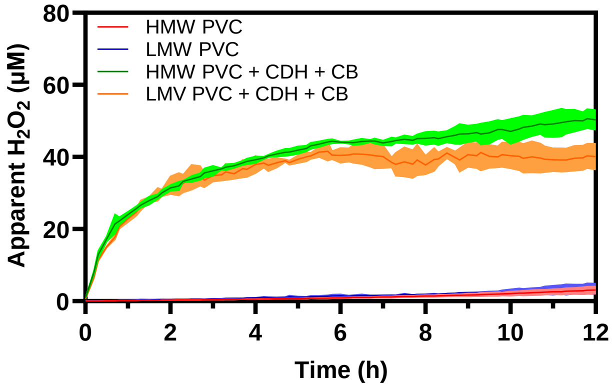


**Figure S2. Apparent $H_2O_2$ generation under reaction conditions used in PVC degradation experiments.** The assays were carried out at room temperature in 50 mM Tris-HCl buffer, pH 8.0, containing 300 mM NaCl, 100 mg/ml PVC, 100 µM Amplex Red and 5 U/ml horseradish peroxidase (HRP). 50 nM cellobiose dehydrogenase (CDH) and 5 mM cellobiose (CB) were present in the reactions to serve as *in situ* hydrogen peroxide source. $H_2O_2$ generation was monitored by following the fluorescence of resorufin. The concentration of hydrogen peroxide was calculated using standard $H_2O_2$ solutions. Note that the hydrogen peroxide generation rates observed in this figure are underestimated due to an increasingly non-linear relationship between the fluorescence and the $H_2O_2$ concentration (due to fluorescence quenching) in the >10 µM range. Error envelopes indicate standard deviation between triplicates.

```
AgPO8          -----------------------------------MATLTQKFHGLLQHPLEPLFLPKND    25
GmPO1          -----------------------------------MSDSKSRLLLFFDRPSEPCFLQKGD    25
GmPO2          -----------------------------------MTDRVKSFQLLFDRPNEPLITPKGE    25
Ceres          --MGRLVLCVL-ALLVGGGIS-DPVKKLQRTVDQTVLDRQYKLLTLFFHPHEPIHIKE--    54
Demetra        MFFNLWFHCNSVTVYFLTEYFILN---NLFAVDPNLVNIQKKVLLLLENWKQVDPDDE--    55
Demetra_new    --MKTVLVLAALIGLVAAGYPLFNNNVKTKTLDPNLVNIQKKVLLLLENWKQVDPDDE--    56

AgPO8          GTLFYDLPERFLTSRYSPIGQNLA--NRFGPNSPASSQVSNDTGVPPTVVTIKDLDELPD    83
GmPO1          DNVAFEVPDHYYPDKYKSLTNTLA--NRFGSGEVRT-------------IPVKNIAL-PD    69
GmPO2          NGAIFQLTQDLLPVDYEDNGIALN--NRFGEEADEK-------------IPLKPLSNPPQ    70
Ceres          ---------------QQEIAASWDLEKNIGLYENAT-------AVHL---------TIQM    83
Demetra        ---------------YYKIGKEYNIEANIESYTNRE-------VVTE---------FLSL    84
Demetra_new    ---------------YYKIGKEYNIEANIEFYTNRE-------VVTE---------FLSL    85

AgPO8          LTFATWIKRRDSFSLFNPEHRKAAGKLTKLFLDQP--NADRLVDVAAYARDRLNAPLFQY    141
GmPO1          LSLPLQLPYNDQFSLFVAKHRRMAGKLIDIFINMR--DVDDLISLCSYCQMRVNPYMFNY    127
GmPO2          FPIASQLPTDADFSLFLPRHQEMATEVIDVLMNIPENQLDDLLSSCVYARGRLNPQLFNY    130
Ceres          LHNNYQVPRGVPFTVLESVHRFEISVYYSLLYSAK--TYDTFYKTAVFLRQHVNENLFVN    141
Demetra        YKTGFTA-KNQIFSIYYENQALEVRALYRLFYYAK--DFETFYKTAAFARVWLNEGQFIY    141
Demetra_new    YKTGFTA-KNQIFSIYYENQALEVRALYRLFYYAK--DFETFYKTAAFARVWLNEGQFIY    142

AgPO8          ALSVALLHRPDTKSVSVPSLLHLFPDQFIDPAAQVRMMEEGSIVLD-ENRMPIP------    194
GmPO1          CLSVAILHRPDTKGLQVPPVVETFPDKFMDPKVFRKARETTTVNASG-NRMPIT------    180
GmPO2          CYSVVLMHRRDTRNVPIQNFAETFPSKFLDSQAFAQARETAAVFPRGIPRTPII------    184
Ceres          VLSVVILHRSDTQDIRIPPIYDVFPSYFHNGEIMTTAQRITTHGQRMLEHYPSTYVW---    198
Demetra        AFYIAVIHRADTRGIVLPAPYEIWPEYFVNSDVLAKINRIQMQKGLILPETAQYYGVLAK    201
Demetra_new    AFYIAVIHRADTRGIVLPAPYEIWPEYFVNSDVLAKINRIQMQKGLILPETAQYYGVLAK    202

AgPO8          ------I---PMNYTATDAEPEQRMAFFREDIGVNLHHWHWHLVYPASGP-PD--VVRKD    242
GmPO1          ------I---PTNYTASNSEPEQRVAYFREDIGINLHHWHWHLVYPFEAA-RE--IVKKD    228
GmPO2          ------I---PRDYTATDLEEEHRLAYWREDIGINLHHWHWHLVYPFTASDRS--IVAKD    233
Ceres          ENN-VVIRHNETAWPYYCNTESMPVSYFTHDVTLNALYYNIKLAYPIWLRSDA--CAIKE    255
Demetra        DNAYYFYANYSGPWTYE--NNENLLSYFIEDVAWNSYYYYFHSKLQFWEKGENAIGPFKE    259
Demetra_new    DNAYYFYANYSGPWTYE--NNENLLSYFIEDVAWNSYYYYFHSKLQFWEKGENAIGPFKE    260

AgPO8          RRGELFYYMHQQLLARYQIDRYAQGLGRIEPLANLREPVREAYYPKLLRTSNNRTFCPRY    302
GmPO1          RRGELFYYMHQQIIARYNAERLCNGLGRVTRYSDFRAPIGEGYFPKLDSQVASRSWPPRF    288
GmPO2          RRGELFFYMHQQIIARYNCERINNSLKRVKKFNNWREPIPEAYFPKLDSLTSSRGWPPRQ    293
Ceres          KRGELFFFWNKQLLARYYMERLSVGLGEIPEL-GLNEVEEGYVSGLLYHN--GIPYPVRP    312
Demetra        RRGEIYYFIYQQILARYYLERLSNGLGEIPRF-NWNDRLQAGYYPLLTTH--QIPFAQRN    316
Demetra_new    RRGEIYYFIYQQILARYYLERLSNGLGEIPRF-NWNDRLQAGYYPLLTTH--QIPFAQRN    317

AgPO8          PGMTISDVARSADRLEVRIADIESWLPRVLEAIDAGFAVSDDGVRVPLDETRGIDVLGNI    362
GmPO1          ANTVIRDIDRPVNEIKIDVFQLETWRDRFLQAIDSNAINMPNGRKVPLNEETGIDELGNL    348
GmPO2          ANMTWQDLNRPVDGLNVTISDMERWRRNLEEAVSMGTVTLPDGSTRPL----DIDTLGNM    349
Ceres          NHLVLNHQTW----HAEAIEEIEVYENRIRDMIDQGFYITNTGEHVSINSPDSIDVLGRL    368
Demetra        GDYYLANDD-----NIEDIQFVDSYEKTFLQFLQKGQFKAYK-QEVDLYNSKSVNFVGNY    370
Demetra_new    GDYYLANDD-----NIEDIQFVDSYEKTFLQFLQKGQFKAYK-QEVDLYNSKSVNFVGNY    371

AgPO8          LERSAIS----INRNLYGDVHNMGHVLLAFIH-----DPRGTYLESSGVMGGVATAMRDP    413
GmPO1          MESSILS----LNRGYYGDLHNMGHVFIAYSH-----DPDHRHLEEYGVMGDSATAMRDP    399
GmPO2          VEASILS----PNRELYGSVHNNGHSFSAYVH-----DPSHRYLENFGVIADEATTMRDP    400
Ceres          IEANVDS----PNVQYYKDFISIWKKVLGNSLVHESVAFNGIPLVVPSVLEQYQTALRDP    424
Demetra        WQANVDLYEKVPQRNYLRSYEDAARRILGAAPR-----NSYENLNVPTALDFYQTSLRDP    425
Demetra_new    WQANVDLYEKVPQRNYLRSYEDAARRILGAAPR-----NSYENLNVPTALDFYQTSLRDP    426

AgPO8          IFYRWHKFIDNIFLRNKA--RLAPYTMAELSNSNVTLEALETQLDRAGGAVNSFVTFWQR    471
GmPO1          VFYRWHAYIDDIFNLYKS--KLTPYGDSQLDYPGIRVSSI--SVEGPAG-ANRFATQWQQ    454
GmPO2          FFYRWHAWVDDLFQKHKESNFVRPYSRSELENPGVQVTSV--SVETQGSPQNVLSTFWMS    458
Ceres          AYYMIMKRVLKLFNLWHE--HLPHYTTKELSVPSVKIEKVE---------VDKLLTYFEY    473
Demetra        AFYQLYAKILDFINQYKE--YLEPYTQDVLHFVGVKINDVK---------VDKLVTYFEY    474
Demetra_new    AFYQLYAKILDFINQYKE--YLEPYTQDVLHFVGVKINDVK---------VDKLVTYFEY    475

AgPO8          SQVDLRAG--------IDFSAAGSAFVSFTHLQCAPFVYRLRINSTARSNRQDTVRIFLL    523
GmPO1          SLVELSQG--------LDFTPRGSVLAKFTHLQHEEFTYVIEVNNTSGQSKMGTFRVFMA    506
GmPO2          SDVDLSRG--------LDFSNRGPVYARFTHLNHRPFRYVIKVNNS-GNARRTTVRIFIS    509
Ceres          TNFNVTNHLHLNEIECNNVINTKSVLVQRTRLNHKVFTVRVNVK--SGVAKHVTVRFFLA    531
Demetra        FDWNATNAVYLSEQQLD--TGSPSYIVRQPRLNNQPFTVTIDIK--SDVESEAVIKIFIG    530
Demetra_new    FDWNATNAVYLSEQQLD--TGSPSYIVRQPRLNNQPFTVTIDIK--SDVESEAVIKIFIG    531
```

```
AgPO8         PRQNEQGRPLSFEDRRLLAIELDSFRVNLRPGMNNIVRQSSNSSV--------TIPFERT      575
GmPO1         PKTDERGQPLAFEDQRRLMIELDKFTRGLKPGNNTIRQRSLDSSV--------TIPFERT      558
GmPO2         PKFDERNLAWSLVDQRKMFIEMDRFVTPLKAGENTITRQSTESTF--------TIPFEQT      561
Ceres         PKYDSVGNEIPLNVNTQNFLLIDIFNYELKEGDNLITRVSSDNLLVTDEIDSASVLFNKV      591
Demetra       PKYDGNGYPIDLENNWVNLVEIDWFTHKLTSGQNKIERKSENFFWFKEDSVSVSKIYE--      588
Demetra_new   PKYDGNGYPIDLENNWVNLVEIDWFTHKLTSGQNKIERKSENFFWFKEDSVSVSKIYE--      589

AgPO8         FGNVEQA-----NAGNAQSRFCGCGWPAHMLLPKGNANGVEFDLFAMVSRFEDDNANVN-      629
GmPO1         FRNQANRPGDPGSATAAEFDFCGCGWPHHMLIPKGTEQGYPVVLYVMVSDWNADKIEQD-      617
GmPO2         FRDLSVQADDPRRVDLAAFNFCGCGWPQHMLVPKGTEAGAPYVLFVMLSNYDLDRIDEPG      621
Ceres         DSALQGHGQYMLNMK-----QNILKTPRHLLLPKGRVGGMPFVLMVYISEYHAPNDVHRG      646
Demetra       ---LLNNGQVPRYMI-----EKFLLLPRRLLLPRGTEGGVPFQFFVFVYPYQAPYKEWEP      640
Demetra_new   ---LLNNGQVPRYMI-----EKFLLLPRRLLLPRGTEGGVPFQFFVFVYPYQAPYKEWEP      641

AgPO8         YDENAGCDDSYAFCGLRDRVYPSRRAMGFPFDRRASNGVRSVADFVAPYKNMRLATVTLR      689
GmPO1         --TVGACNDAASYCGLRDRKYPDKRHMGFPFDRRSE--ARNLTDFLKP--NMATRDCTIK      671
GmPO2         NSPEISCKEASSFCGLRDRKYPDKRAMGFPFDRPSR-TATSIEDFILP--NMALQDITIR      678
Ceres         TVETST---------IDNTIRLTSDTLGFPVDRPLFP------WMLTGVENIFLQDVQIY      691
Demetra       MK----------------EFVVDNKPFGYPFDRPVTE------SYYFTQPNMYFKDVYIY      678
Demetra_new   MK----------------EFVVDNKPFGYPFDRPVTE------SYYFTQPNMYFKDVYIY      679

AgPO8         FMNTIIDRPT---------N-----      700
GmPO1         FTDAIREGTQ---------RQ----      683
GmPO2         LNNVVEANPR---------NP--RT      692
Ceres         HKPTTEVTGVPVYVE----------      706
Demetra       QEGEE----YPYYTSYWSQNQVPKH      699
Demetra_new   QEGEE----YPYYTSYWSQNQVPKH      700
```

**Figure S3. Sequence alignment of two *Galleria mellonella* hexamerins ("Ceres" and "Demetra") compared to three insect phenoloxidases (POs).** *Ag*PO8, PO from *Anopheles gambiae* (NCBI accession number XP_315074.1); *Gm*PO1 and *Gm*PO2, POs from *Galleria mellonella* (NCBI accession numbers XP_052751666.1 and XP_026755063.2, respectively); Ceres and Demetra, hexamerins from *Galleria mellonella* previously reported by Sanluis-Verdes et al. [1] (NCBI accession numbers XP_026756459.1 and XP_026756396.1, respectively); Demetra_new, most recent variant of "Demetra" sequence (NCBI accession number XP_052756923.1) which is found in the new version of the *Galleria mellonella* whole genome (NCBI accession number NW_026442147.1). Red color indicates histidine residues forming a bi-nuclear copper-binding cluster (the active site) in phenoloxidases. Note that all of these conserved histidines are lacking in "Ceres" and most of them (5 out of 6) are lacking in "Demetra", indicating the loss of catalytic activity. The active site residues were annotated using the X-ray structure of *Ag*PO8 showing copper atoms (PDB: 4YZW). Yellow color indicates the putative Sec/SPI signal peptide observed in "Ceres" and in the new version of "Demetra" sequence, but not in original "Demetra" or any of the POs. Signal peptide prediction was carried out using SignalP 6.0 (https://services.healthtech.dtu.dk/services/SignalP-6.0/). Multiple sequence alignment was performed with Clustal Omega (https://www.ebi.ac.uk/jdispatcher/msa/clustalo).

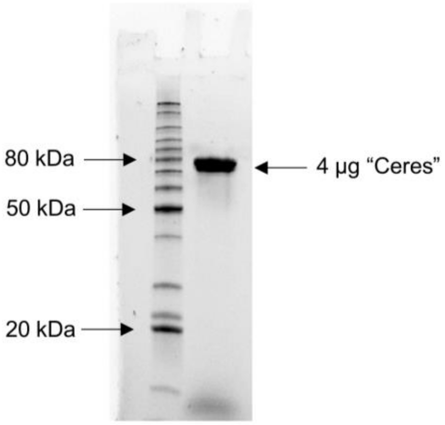


**Figure S4. SDS-PAGE of “Ceres” after purification.** The predicted molecular weight of the mature protein with the affinity tag (not accounting for any potential post-translational modifications) is 80.7 kDa. The molecular weight calculation was carried out using ProtParam tool (https://web.expasy.org/protparam/).

1 DPVKKLQR**TV DQTVLDR**QYK **LLTLFFHPHE PIHIKEQQEI AASWDLEK**NI
51 GLYENATAVH LTIQMLHNNY QVPR**GVPFTV LESVHR**FEIS VYYSLLYSAK
101 **TYDTFYK**TAV FLR**QHVNENL FVNVLSVVIL HRSDTQDIRI PPIYDVFPSY**
151 **FHNGEIMTTA QRITTHGQRM LEHYPSTYVW ENNVVIR**HNE TAWPYYCNTE
201 SMPVSYFTHD VTLNALYYNI K**LAYPIWLRS DACAIK**EK**RG ELFFFWNK**QL
251 LARYYMERLS VGLGEIPELG LNEVEEGYVS GLLYHNGIPY PVRPNHLVLN
301 HQTWHAEAIE EIEVYENRIR DMIDQGFYIT NTGEHVSINS PDSIDVLGR**L**
351 **IEANVDSPNV QYYKDFISIW K**KVLGNSLVH ESVAFNGIPL VVPSVLEQYQ
401 TALR**DPAYYM IMKR**VLK**LFN LWHEHLPHYT TKELSVPSVK IEKVEVDK**LL
451 TYFEYTNFNV TNHLHLNEIE CNNVINTKSV LVQRTR**LNHK VFTVR**VNVK**S**
501 **GVAKHVTVR**F FLAPKYDSVG NEIPLNVNTQ NFLLIDIFNY ELK**EGDNLIT**
551 **RVSSDNLLVT DEIDSASVLF NKVDSALQGH GQYMLNMKQN ILK**TPR**HLLL**
601 **PKGR**VGGMPF VLMVYISEYH APNDVHR**GTV ETSTIDNTIR LTSDTLGFPV**
651 **DR**PLFPWMLT GVENIFLQDV QIYHKPTTEV TGVPVYVEAH HHHHH

**Figure S5. LC-MS/MS analysis of the “Ceres” sample.** The figure shows the protein sequence of mature “Ceres” with a C-terminal affinity tag (“-AHHHHHH”). Red color indicates the parts of the sequence that were observed in peptides resulting from tryptic cleavage of "Ceres”. This represents a sequence coverage of 46%.

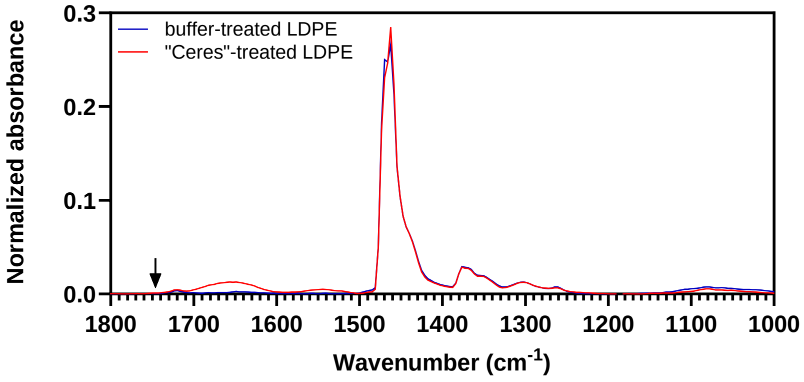


**Figure S6. FTIR analysis of LDPE film treated with "Ceres" by applying multiple drops of a protein solution on the surface.** The figure shows parts of the FTIR spectra of "Ceres"-treated film and buffer-treated film after 14 applications of 5 µl of 20 mM HEPES, pH 7.0, supplied with 150 mM NaCl and 5% (v/v) glycerol, and containing or lacking 1 mg/mL "Ceres". The experiments were carried out at room temperature for 3 days using additive-free LDPE films, produced in-house. The droplets were applied at the same area of the films. Each trace shown in the figure represents the average signal of 3 independent measurements of the same film sample. The black arrow indicates the expected position (≈1,745 $cm^{-1}$) of the carbonyl peak which was not observed in our experiments but was previously reported and taken to show polyethylene oxidation by "Ceres". The absorbance signals were normalized using the $CH_2$ asymmetric C–H stretch peak of PE (≈2,913 $cm^{-1}$) as the reference (not visible in this zoomed-in view). The data were acquired with 8 $cm^{-1}$ resolution after washing the films with water and EtOH.

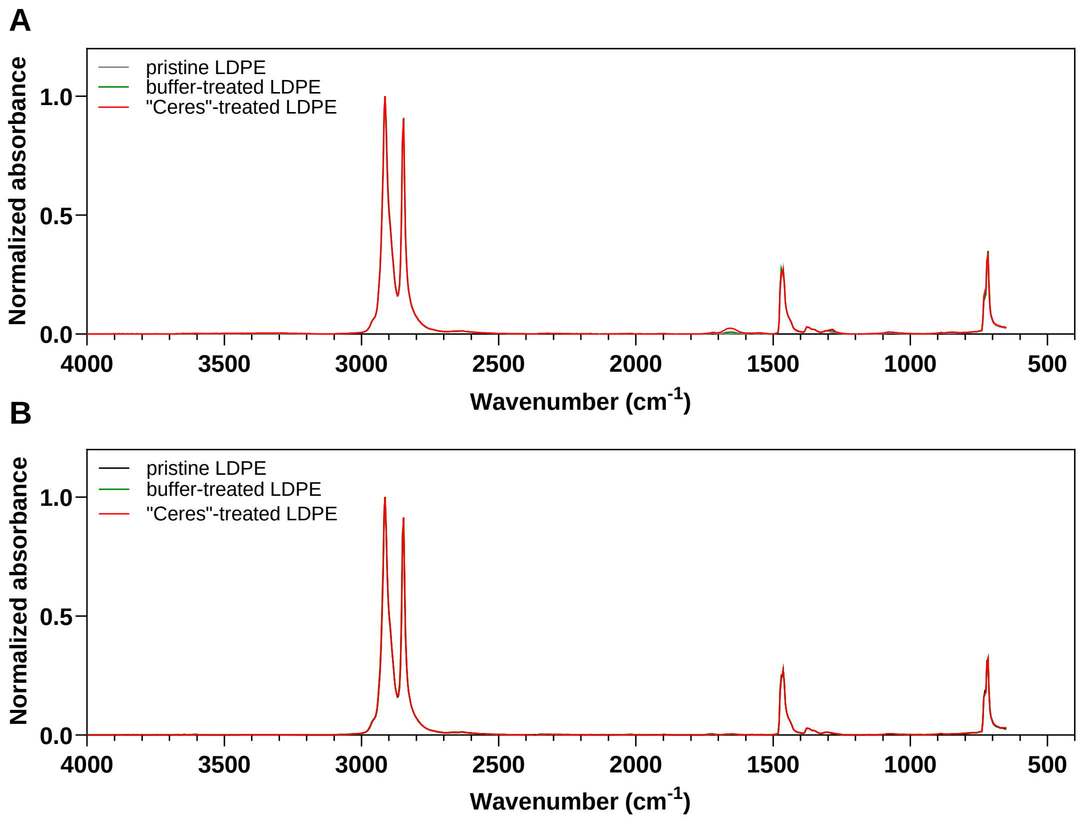


**Figure S7. FTIR spectra of pristine, buffer-treated and “Ceres”-treated LDPE films before (panel A) and after cleaning with 1 M NaOH (panel B).** These data were acquired with 8 $cm^{-1}$ resolution and used to produce Fig. 5 (see the main manuscript for the reaction conditions). The absorbance signals were normalized using the $CH_2$ asymmetric C–H stretch peak of PE (≈2,913 $cm^{-1}$) as the reference. Each trace shown in the figure represents the average of signals obtained with 3 film discs. Note that the traces are overlapping almost completely.

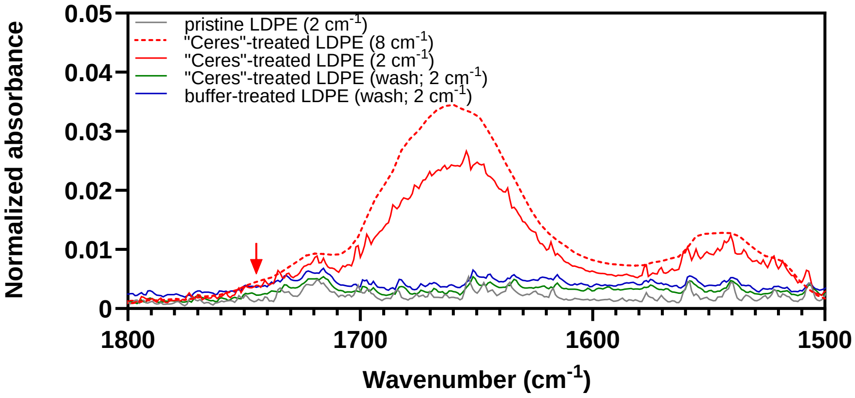


**Figure S8. FTIR spectra of "Ceres"-treated additive-free LDPE film before and after washing with 1 M NaOH.** LDPE film discs were incubated at 25 °C for 70 hours in 20 mM HEPES, pH 7.0, supplied with 150 mM NaCl and 5% (v/v) glycerol, in the presence or absence of 3 mg/mL "Ceres". Red arrow indicates the expected position (≈1,745 $cm^{-1}$) of the carbonyl peak which was not observed in our experiments but was previously reported and taken to show polyethene oxidation by "Ceres". Each trace shown in the figure represents the average of signals obtained with 2 film discs. The absorbance signals were normalized using the $CH_2$ asymmetric C–H stretch peak of PE (≈2,913 $cm^{-1}$) as the reference (not visible in this zoomed-in view). The data were acquired with 2 $cm^{-1}$ resolution or with 8 $cm^{-1}$ resolution as indicated in the figure legend.

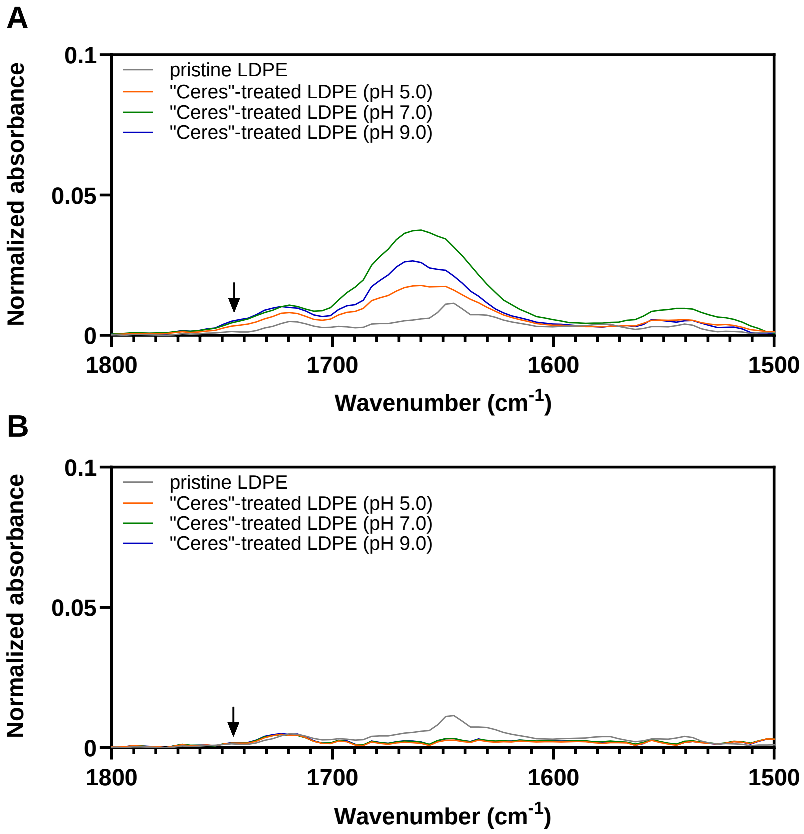


**Figure S9. FTIR spectra of a pristine and "Ceres"-treated commercial LDPE film before (panel A) and after cleaning with 1 M NaOH (panel B).** LDPE film samples were incubated at 25 °C for 70 hours in reaction buffer containing or lacking 1 mg/mL "Ceres". Three reactions buffers were used, to assess potential pH effects: (1) 50 mM sodium acetate, pH 5.0; (2) 50 mM HEPES, pH 7.0; (3) 50 mM sodium phosphate, pH 9.0. Black arrows indicate the expected position (≈1,745 $cm^{-1}$) of the carbonyl peak which was not observed in our experiments but was previously reported and taken to show polyethene oxidation by "Ceres". Each trace shown in the figure represents the average of signals obtained with 3 film discs. The absorbance signals were normalized using the $CH_2$ asymmetric C–H stretch peak of PE (≈2,913 $cm^{-1}$) as the reference (not visible in this zoomed-in view). The data were acquired with 8 $cm^{-1}$ resolution.

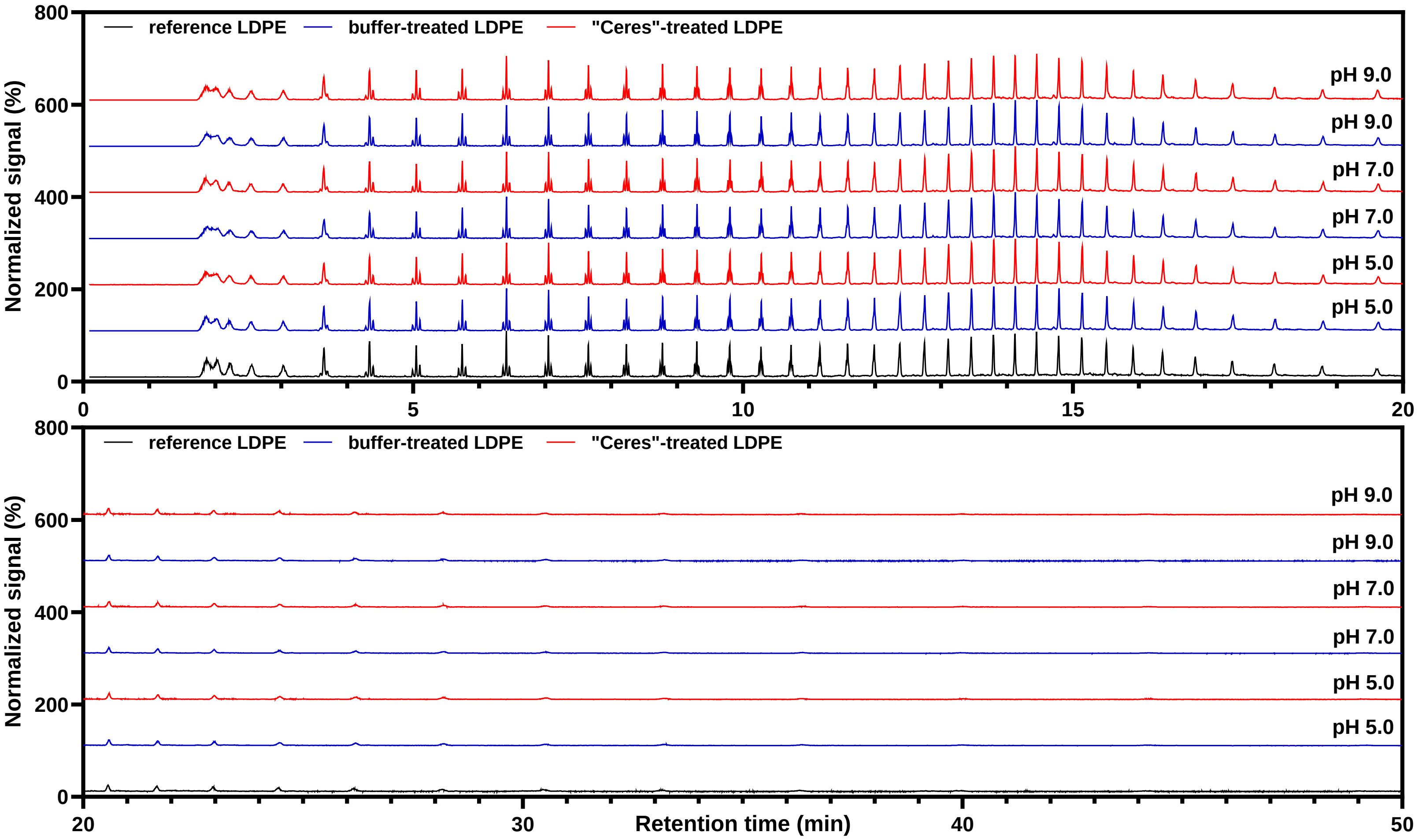


**Figure S10. Pyrolysis gas chromatography/mass spectrometry analysis of a buffer-treated and "Ceres"-treated commercial LDPE film.** LDPE film samples were incubated at 25 °C for 70 hours in reaction buffers containing or lacking 1 mg/mL "Ceres". Three reactions buffers were used to assess the potential pH effects: (1) 50 mM sodium acetate, pH 5.0; (2) 50 mM HEPES, pH 7.0; (3) 50 mM sodium phosphate, pH 9.0. Note that the treatment of LDPE with "Ceres" did not result in the formation of novel compounds released after thermal degradation of materials. The chromatograms were plotted using total ion count (TIC) signal of the mass selective detector. A commercial untreated LDPE film (grade 22H594; INEOS Olefins & Polymers Europe, Rolle, Switzerland) was used as a reference PE sample.
